# Systematic tools for reprogramming plant gene expression in a simple model, *Marchantia polymorpha*

**DOI:** 10.1101/2020.02.29.971002

**Authors:** Susanna Sauret-Güeto, Eftychios Frangedakis, Linda Silvestri, Marius Rebmann, Marta Tomaselli, Kasey Markel, Mihails Delmans, Anthony West, Nicola J. Patron, Jim Haseloff

## Abstract

We present the OpenPlant toolkit, a set of interlinked resources and techniques to develop Marchantia as testbed for bioengineering in plants. Marchantia is a liverwort, a simple plant with an open form of development that allows direct visualization of gene expression and dynamics of cellular growth in living tissues. We describe new techniques for simple and efficient axenic propagation and maintenance of Marchantia lines with no requirement for glasshouse facilities. Marchantia plants spontaneously produce clonal propagules within a few weeks of regeneration, and lines can be amplified million-fold in a single generation by induction of the sexual phase of growth, crossing and harvesting of progeny spores. The plant has a simple morphology and genome with reduced gene redundancy, and the dominant phase of its life cycle is haploid, making genetic analysis easier. We have built robust Loop assembly vector systems for nuclear and chloroplast transformation and genome editing. These have provided the basis for building and testing a modular library of standardized DNA elements with highly desirable properties. We have screened transcriptomic data to identify a range of candidate genes, extracted putative promoter sequences, and tested them *in vivo* to identify new constitutive promoter elements. The resources have been combined into a toolkit for plant bioengineering that is accessible for laboratories without access to traditional facilities for plant biology research. The toolkit is being made available under the terms of the OpenMTA and will facilitate the establishment of common standards and the use of this simple plant as testbed for synthetic biology.

## INTRODUCTION

Synthetic biology is a field that aims to bring formal engineering approaches to the design, construction and analysis of biological systems ^1^. Most recent advances in synthetic biology have used microbes as a platform. Transferring this approach to plants is challenging due to our limited understanding of plant genetic networks and increased complexity due to multicellularity ^2^. However, plants harbor great potential for synthetic biology ^3^. Synthetic biology approaches can enable the production of compounds in specific organelles and cell types, useful for food, medicine or industry. In plants, much energy production and biosynthetic capacity resides within chloroplasts. These photosynthetic organelles have a simple and prokaryote-like genome, with the ability to support high levels of transgene expression and metabolic activity. The proper harnessing of plants’ capacity for bioproduction will require access to tools that allow multi-scale manipulation of the nuclear and chloroplast genomes, building of synthetic metabolic pathways and engineering of existing or new harvestable organs on the plant. A major benefit is that engineered plants can potentially be grown at large scale, using globally accessible and low-cost agricultural production methods.

Much of our current knowledge of plant genetics and genome function comes from study of model angiosperm weeds and crops, such as Arabidopsis, maize, rice, tomato and tobacco. Work with these plants is beset with one or more difficulties due to relatively long generation times, inefficient transformation and slow regeneration, functional redundancy in diploid genomes and difficult access to early stages of growth. A simple prototype system is clearly needed. *Marchantia polymorph*a is the best-studied species of liverwort, an early divergent land plant lineage ^4^. Marchantia is proving to be a powerful and versatile experimental system for plant biology studies ^5–7^ and an emerging platform for plant synthetic biology ^8^. Marchantia has a small size and grows fast and resiliently under laboratory conditions. In contrast to higher plant systems, the dominant phase of its life cycle is haploid, with distinct male and female plants. It can be propagated both by spores derived from sexual reproduction and also through asexual clonal propagules called gemmae (Figure 1A). In just a week, both germinating spores and gemmae can give rise to tiny plantlets with a morphologically simple, sheet-like and modular plant body called thallus. Thallus grows from apical stem cells located in invaginated notches at the thallus extremities ^9^. The early stages of development in Marchantia happen in an open fashion, allowing easy and direct observation of formative processes during morphogenesis in a way amenable to rapid and high-throughput analysis. The morphological simplicity of Marchantia is matched by a small genome size (220 Mb) with low genetic redundancy ^7^. It shares most of the characterized gene regulatory components of more recently diverged land plants like angiosperms, but gene families are often sparsely populated, and haploidy makes genetic analysis easier ^7^. Altogether, Marchantia has the potential to be a unique multicellular testbed to study and engineer the links between genetic networks, cell processes, cell communication, cell fate decisions and tissue-wide physical processes that drive morphogenesis ^5^.

**Figure 1.**
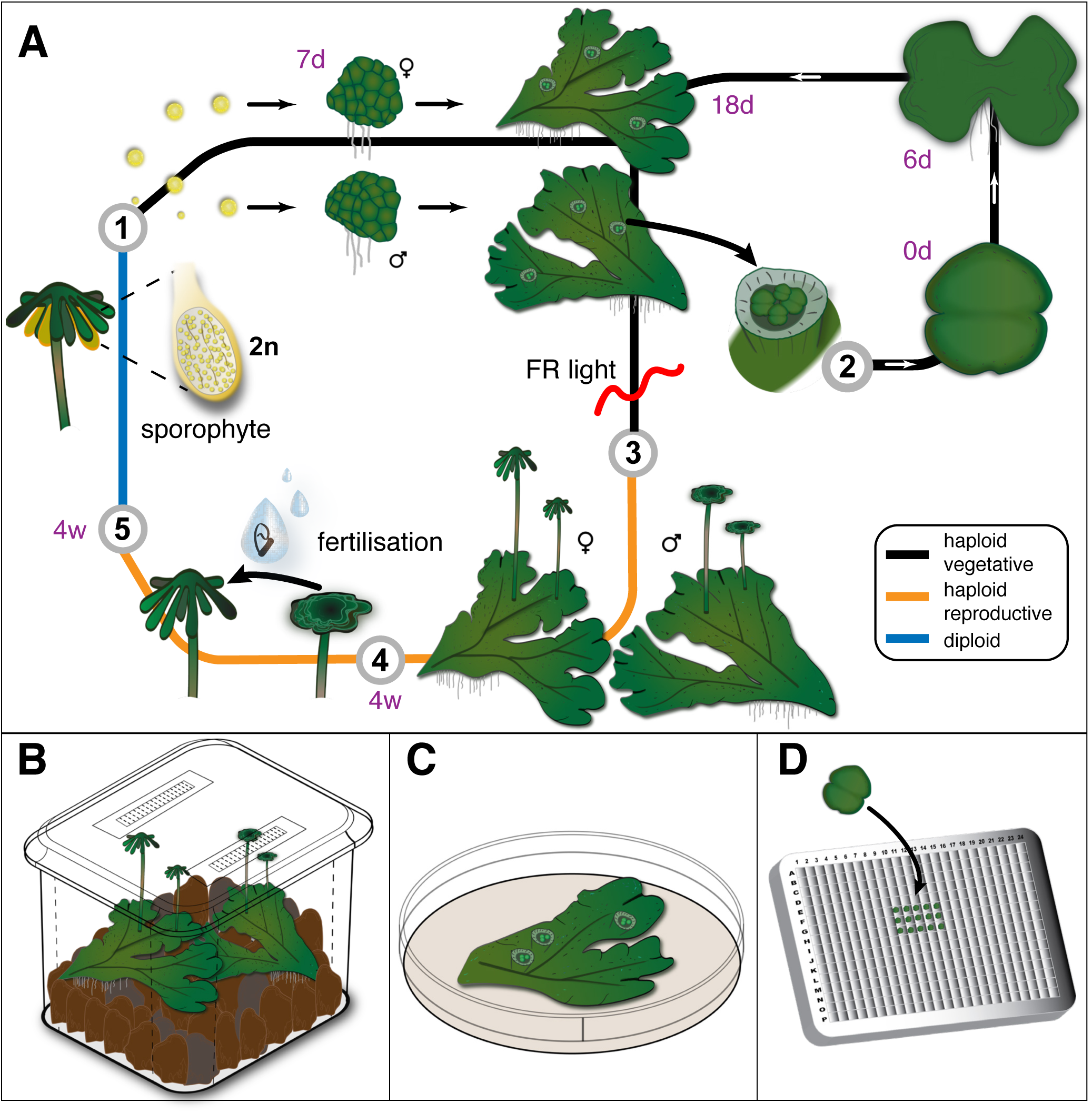
Use of the Marchantia life cycle for propagation in the laboratory. (A) Marchantia has two stages in its life cycle. As a bryophyte, the haploid (n) gametophyte form of the plant is dominant, larger, and longer-lived (stages 1-4) than the transient diploid (2n) sporophyte (stage 5). Marchantia is dioecious with separate male and female plants. The life cycle is shown starting with the germination of a mixture of male and female haploid spores (stage 1). These germinate to form male and female sporelings and plantlets after 7 days, which grow into similar-looking plants with a prostrate thallus. Asexual reproduction occurs via formation of clonal propagules called gemmae, which are produced spontaneously in cup-like structures on the dorsal side of both male and female thalli (stage 2). A gemma can be removed, germinated, and will develop into a plant with new gemmae cups in approximately 3 weeks. This provides a simple method for fast clonal propagation of lines. Vegetative tissues can be exposed to far red (FR, wavelength 730-740 nm) light in order to trigger the sexual phase of the life cycle (stage 3). Different reproductive organs develop on male and female plants, and mature within 4 weeks (w). Motile sperm cells are produced by male reproductive organs (antheridia) and can be transferred onto female reproductive organs (archegonia), where they can fertilize an egg cell (4). The resultant diploid zygote develops as a sporophyte (stage 5) and will produce a capsule (sporangia) with a large quantity of haploid spores (stage 1) after 4 weeks. (B) Marchantia can be propagated axenically throughout its life cycle for production of spores using Microboxes, with depth-filtration for high efficiency gas exchange and minimal water loss. (C) Petri-dishes can be used for amplification of plantlets and gemmae propagation. (D) Gemmae can be cultured for microscopic observation in multiwell plates, allowing rapid screening of lines.

The implementation of synthetic biology approaches in plants calls for rapid and efficient methods for assembly of DNA circuits. Golden Gate Cloning methods (such as MoClo and GoldenBraid ^10, 11^) are based on Type IIS restriction enzymes ^12^ and enable fast generation of complex DNA circuits from standardized basic DNA parts (e.g., promoters, CDS and terminators). Golden Gate Cloning based toolkits have already been developed for other model plants ^13, 14^. We have established a common syntax for modular DNA parts ^15^ and developed a simplified recursive Type IIS assembly method, the Loop assembly ^16^.

In this paper, we describe work that exploits these technical advances in DNA assembly and seek to improve and extend the set of compatible tools to develop an integrated system for reprogramming Marchantia and exploit its unique characteristics as a testbed for synthetic biology experiments in plants. (i) We adopted techniques to allow for axenic propagation of Marchantia throughout the life cycle, (ii) improved the throughput of Agrobacterium-mediated nuclear genome transformation of sporelings, and (iii) developed a simplified and more efficient technique for generation of homoplasmic plastid transformants. (iv) We swapped the plasmid backbone of Loop assembly vectors to provide higher yields of plasmid DNA, (v) expanded the repertoire of vectors to allow chloroplast transformation and (vi) delivery of *Cas9* genes and guide sequences for efficient nuclear gene editing. (vii) We constructed a standardized and modular library of DNA parts for expression in Marchantia. These include antibiotic resistance genes for selection of successful transformants, signal peptides, and fluorescent proteins for multispectral imaging. (viii) We established higher throughput methods for screening transformed Marchantia plants using multiwell plates and fluorescence microscopy. Using this new suite of techniques, (ix) we surveyed the Marchantia genome for transcriptionally active genes, synthesized proximal promoter sequences, domesticated these as standardized DNA parts for Loop assembly, and constructed and tested reporter gene fusions *in vivo*. (x) We have created a novel set of compact gene elements with highly desirable properties, such as promoters that confer ubiquitous and specific patterns of gene expression. The increased scale of Marchantia experiments together with the need for standardized practices to support reproducibility, have demanded the establishment of research data management frameworks, and we have adopted the use of Benchling. Benchling provides a laboratory information management systems (LIMS), electronic notebook, molecular biology suite and collaboration platform, and enables, for instance, a shared curated registry of annotated DNA parts and linked characterization data. (xi) To support access to the wider community, the OpenPlant toolkit sequences and metadata are shared through a Benchling public folder. (xii) We have also deposited plasmids in Addgene for global distribution under an OpenMTA license ^17^ for public sharing and open access. The OpenMTA facilitates open exchange of biological materials, and was designed to support openness, sharing and innovation in global biotechnology. In contrast to the widely used UBMTA, the OpenMTA allows redistribution and commercial use of materials, while still acknowledging the creators and promoting safe practices. (xiii) We describe and distribute the relevant protocols at protocols.io/OpenPlant. The establishment of this set of open, interlinked resources and techniques for Marchantia is intended to encourage the establishment of a shared commons to facilitate scientific exchange and innovation in advanced bioengineering with this new testbed for plant synthetic biology.

## RESULTS AND DISCUSSION

### Axenic culture for spore production

Certain specialized features of Marchantia life cycle provide substantial advantages to the experimental scientist. We have adopted techniques to allow axenic culture of Marchantia throughout its life cycle (Figure 1). The haploid gametophyte form of the plant is dominant, larger, and longer-lived than the transient diploid sporophyte stage (Figure 1A). The gametophyte takes the form of a flattened thallus with rhizoids emerging from the ventral surface, with photosynthetic chambers and other specialized cellular structures forming on the dorsal surface (Figure 1A-1).

Marchantia can reproduce both asexually and sexually. Asexual reproduction occurs via clonal propagules called gemmae, which are formed inside conical splash cups (Figure 1A-2). Each gemma is derived from a single epidermal cell at the base of a cup that expands and divides to form a group of cells carried on a short stalk. The cells continue to proliferate in regular fashion to form a bilobed gemma with a conserved morphology. This is eventually detached from the stalk, and can be dispersed from the cup, typically by water splash. Gemmae can be used for simple vegetative propagation and amplification of plantlets during experiments, grow well in Petri dishes (Figure 1C) and can be stored in the cold or cryopreserved when long term storage is necessary. We have simplified a previously published cryopreservation method to avoid the need for cryoplates, making it a more routine process ^18^ (Figure S4 and Materials and Methods). Gemmae grow in exposed fashion, allowing easy access to morphogenetic processes, and we have adapted multiwell plates for culture and screening of lines under microscopic observation (Figure 1D, S16, and Materials and Methods).

Upon induction of formation of male and female reproductive organs (gametophores) and sexual reproduction, each cross can result in the formation of millions of single cell spores ^5^ (Figure 1A-1). Efficient nuclear and chloroplast transformation methods both rely on the use of sterile sporelings (germinating spores) as a target tissue. In order to allow reliable production of large quantities of sterile Marchantia spores, we have adopted the use of Microbox micropropagation containers with a specially designed lid that allows high gas exchange, limited dehydration and blocks the entry of contaminants such as bacteria and fungi into the culture (Figure 1B, S1 and Materials and Methods). Plantlets or gemmae grown under axenic conditions (Figure 1C) can be used to inoculate the containers and grown in a growth room or controlled environment cabinet with limited need for watering or other maintenance. Supplementary far red light triggers the sexual phase, with efficient formation of male and female gametophores after 4 weeks in Cam-1 and Cam-2 strains of *M. polymorpha* (Figure S1). In our experience, the widely used Tak-1 and Tak-2 isolates can be slower to respond to far-red light induction. The Cam-1 and Cam-2 isolates have not been extensively back-crossed yet, so a small degree of genetic variability will be introduced after cross fertilization, but no phenotypic variation is evident. Mature sperm can be harvested from antheridia (male sex organs), diluted and transferred to archegonia (female sex organs) (Figure 1A-4, S1). Yellow colored sporangia containing huge numbers of spores, emerge after a further 4 weeks (Figure 1A-5), and can be stored in a cold, desiccated state (Figure S1 and Materials and Methods) and sterilized (Materials and Methods) prior to plating on suitable medium for germination.

### Transformation of sporelings

We have improved the throughput of Agrobacterium-mediated nuclear genome transformation of sporelings ^19^ by scaling down previous protocols in order to use multiwell plates instead of flasks. This allows to easily perform multiple transformations in one experiment, eliminating the transformation step as a bottleneck for the production of lines (Figure S2 and Materials and Methods). Positive transformants are obtained in about two weeks (Figure S2J). The OpenPlant toolkit contains three antibiotic resistance genes commonly used as selectable markers for Marchantia ^20^ : the hygromycin phosphotransferase gene (*hptII*), the modified acetolactate synthase gene (*mALS*) and the neomycin phosphotransferase II gene (*nptII*), that confer hygromycin, chlorsulfuron and kanamycin resistance, respectively (Figure 8).

Chloroplast transformation of Marchantia sporelings is achieved through direct biolistic delivery of DNA-loaded microcarriers into the plastids. DNA constructs contain flanking homologous sequences, and are integrated into a specific chloroplast genome location via homologous recombination ^21^. Previously published chloroplast transformation methods in Marchantia have required the bombardment of a large number of samples to generate a small number of transformed plants ^22^. We have modified the protocol by adopting nanoparticles called DNAdel^TM^ as plasmid DNA carriers (Figure S3 and Materials and Methods). This reduces the time and labor required for loading of the plasmid DNA onto the microcarrier and increases efficiency and reproducibility. Marchantia sporelings regenerate after biolistic delivery without need for hormone treatment, and typically we can use the spores of a single sporangium to obtain about 10 transplastomic plants after eight weeks of selection on antibiotic-containing growth media (Figure S3). PCR analysis of transformants in a typical experiment (n=30) showed that greater than 90% of plants were homoplasmic (Figure S10). The OpenPlant kit includes the aminoglycoside adenyltransferase (*aadA*) resistance gene which confers spectinomycin resistance ^22^ (Figure 8).

### DNA part syntax and domestication

The OpenPlant consortium initiated the establishment of the common genetic syntax with wide support from the international plant science community ^15^ and we have adopted this convention for the composition (and exchange) of plant DNA parts (Phytobricks). The common syntax defines specific 5’ end and 3’ end fusion sites, of four nucleotides in length, for each type of standardized DNA part (called Level 0 parts, L0) (Figure 2A). In this way, it facilitates community-based efforts to build and exchange collections of L0 parts. Examples of different types of L0 parts are: promoter with 5’ UTR (PROM5), coding sequence with start and stop codons (CDS), or 3’ UTR with terminator (3TERM). The 5’ end and 3’ fusion sites on each L0 part enables directional assembly of multiple L0 parts (Figure 2A, e.g. ggag-PROM5-aatg-CDS-gctt-3TERM-cgct). We provide all L0 DNA parts cloned into a new universal acceptor plasmid called pUAP4 (Figure 2B) which is a derivative of the pUAP1, an OpenMTA vector used for cloning of L0 parts in the MoClo kit ^13^. pUAP4 is designed for Loop assembly, with *Sap*I divergent sites flanking the L0 DNA part insertion site, in contrast to the *Bbs*I sites in pUAP1 (Figure S5). Also, instead of the red fluorescence protein (RFP) as a marker for screening of colonies, the pUAP4 encodes LacZ activity, which allows for blue-white screening and is more visible by naked eye than the RFP (Figure S5). The pUAP4 vector also has convergent *Bsa*I sites to assemble multiple L0 parts into a transcription unit using *Bsa*I (Figure 2C and S7). When a L0 part has internal *Bsa*I or *Sap*I sites, it is necessary to domesticate the DNA part (to remove restriction endonuclease recognition sites that would otherwise interfere with the cloning procedure), which can be done together with cloning the part into pUAP4 (Figure S6).

**Figure 2.**
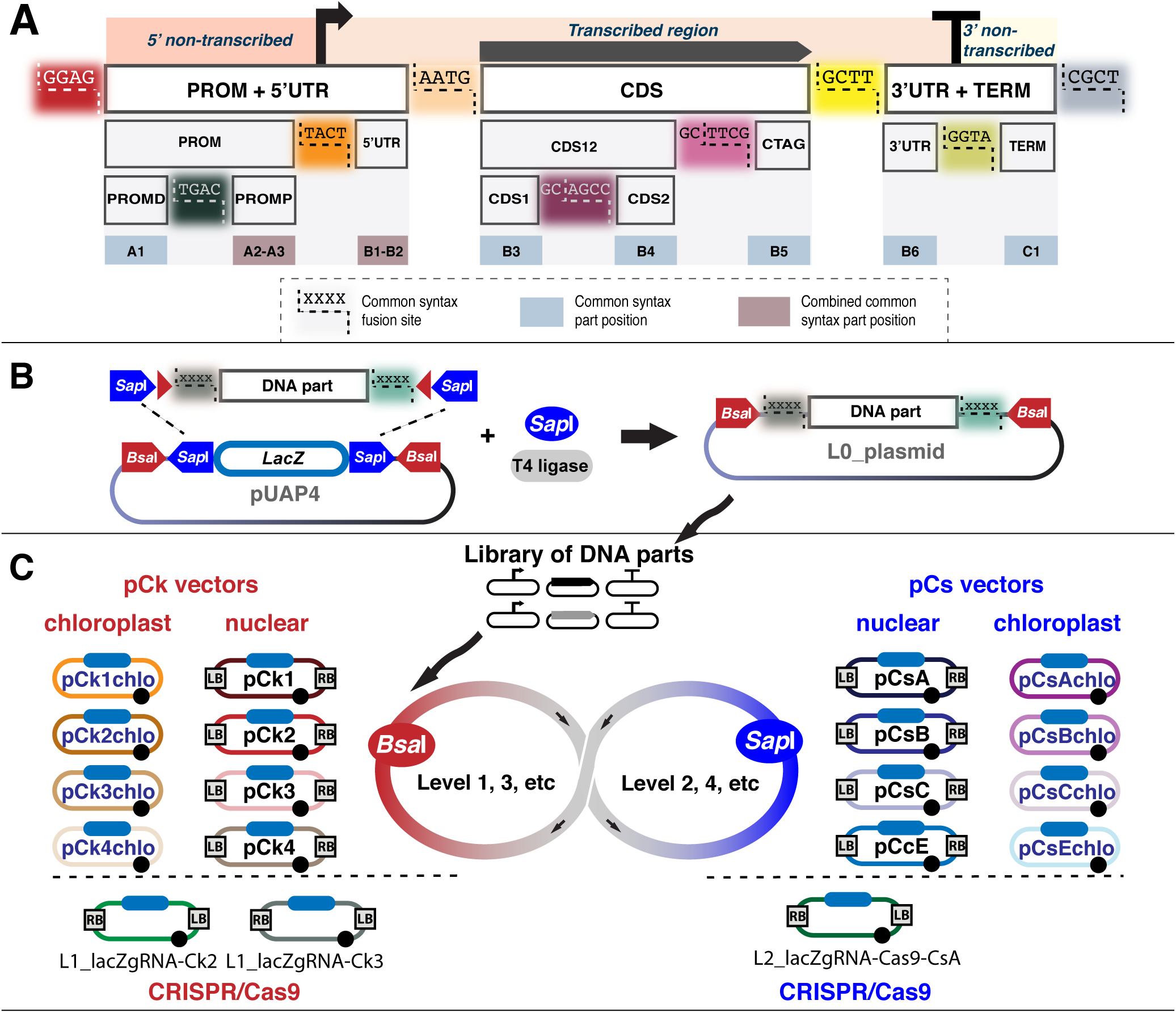
Key elements in the OpenPlant Loop assembly toolkit. (A) The Phytobrick common syntax defines ten DNA part positions and 12 DNA part fusion sites. For Marchantia, we commonly use eight positions and nine fusion sites, by combining positions A2-A3 for proximal promoter (PROMP), and B1-B2 for 5’ untranslated region (5UTR). The other types of parts are: A1 for distal promoter (PROMD), B3 for coding sequence with start codon and no stop codon (CDS1), B4 for coding sequence without start or stop codon (CDS2), B5 coding sequence without start codon and with stop codon (CTAG), B6 for 3’ untranslated region (3UTR) and C1 for transcription terminator (TERM). Parts can span multiple fusion sites, like A1-A3 for promoter (PROM), A1-B2 for promoter with 5’ UTR (PROM5), B3-B4 for coding sequence with start codon and no stop codon for N-terminal fusion with CTAG (CDS12), B3-B6 for coding sequence with start and stop codons (CDS), or B6-C1 for 3’ UTR with terminator (3TERM). (B) Schematic representation of the pUAP4 vector with divergent *Sap*I sites to accept L0 parts and convergent *Bsa*I sites to assemble L0 parts into transcription units (L1). (C) Summary of the Loop acceptor vectors of the OpenPlant toolkit. For nuclear genome transformation: pCk (1,2,3,4) can be used for assembly of L0 parts into a Level 1 plasmid using *Bsa*I, and pCs (A,B,C,E) can be used for assembly of up to four Level 1 plasmids into a Level 2 construct using *Sap*I. The vectors pCkchlo (1,2,3,4) and pCschlo (A,B,C,E) can be used for chloroplast applications. The vectors L1_lacZgRNA-Ck2, L1_lacZgRNA-Ck3 and L2_lacZgRNA-Cas9 are designed for CRISPR/Cas9 genome editing. LB and RB: left and right border repeats respectively from nopaline C58 T-DNA. Filled blue rounded rectangle: lacZα cassette for blue-white screening. Filled black circles: pSa origin or replication.

The OpenPlant toolkit is composed of a set of 50 L0 parts: 11 promoters and 5UTR parts (PROM5=PROM+5UTR, PROM and 5UTRs), 33 CDS-type parts (CDS, CDS1, CDS2, CDS12, CTAG), and 6 3UTR and terminators (3TERM=3UTR+TERM) (Figure 8). L0 DNA parts were either amplified by PCR and cloned, or synthesized into the pUAP4 vector. All L0 parts have been verified by sequencing.

### Recursive assembly of gene parts

Pollak *et al*. ^16^ previously described a method for gene assembly based on recursive use of two Type IIS restriction endonucleases, *Sap*I and *Bsa*I, called Loop assembly. Loop assembly allows rapid and efficient production of large DNA constructs for nuclear DNA transformation and is compatible with widely used standardized DNA parts such as Phytobricks. In a single step reaction with *Bsa*I and T4 ligase, L0 parts can be assembled into a transcription unit (Level 1, L1) (Figure 2C, S7). Then L1 constructs can be assembled together into DNA assemblies with up to four transcription units (Level 2, L2), in a single step reaction with *Sap*I and T4 ligase (Figure S8). The recursive nature of Loop assembly means that four L2 devices can be digested with *Bsa*I and ligated to combine up to sixteen transcription units in a Level 3 device, and so on (Figure 2C). Loop assembly is well suited to automation and we developed methods for assembly using an acoustic-focusing non-contact liquid handling robot, which increases the speed and scale of assembly, while allowing reactions to be performed in sub microliter volumes to reduce cost. We have scaled down both L1 and L2 assemblies to 500 nL using the Labcyte Echo acoustic focusing robot (Materials and Methods).

### Modified vector systems

The manuscript describes an extension of the Loop vector system ^16^ (Figure 2C). The plasmid backbones of the original Loop vectors were based on pGREEN II ^23, 24^, which contains ColE1 and pSA origins of replication. We have replaced the plasmid backbone with that from pCambia/pPZP ^25^. The 7.5 kb pCambia backbone encodes a VS1 origin, the pVS1 RepA replication protein, and StaA protein that confer plasmid stability and higher yields in *E. coli* and stable maintenance in Agrobacterium. All of the modified Loop vectors used for Agrobacterium mediated transformation of the nuclear genome contain the right and left border repeats from nopaline C58 T-DNA (RB and LB) derived from pCambia, and show similar efficiencies of transformation to other plasmids based on this well-tested backbone. The new odd (L1, L3,…) acceptor vectors are collectively called pCk vectors (pCk1, pCk2, pCk3 and pCk4), and the new even (L2, L4,…) acceptor vectors are collectively called pCs vectors (pCsA, pCsB, pCsC and pCsE) (Figure S9A&B). We have not observed untoward toxic effects due to the increase in plasmid copy number, and the new plasmids are easier to harvest in useful quantities for routine cloning operations, without the need to increase the scale of host microbial cultures.

We have expanded the repertoire of vectors to include not only wide-host range plasmids for Agrobacterium-mediated nuclear transformation, but also vectors for chloroplast transformation via biolistic DNA delivery, and vectors for CRISPR-Cas9 mediated genome editing (Figure 2C).

### DNA tools for chloroplast genome modification

The Loop set of vectors for chloroplast transformation are collectively called pCkchlo (pCk1chlo, pCk2chlo, pCk3chlo and pCk4chlo) and pCschlo (pCsAchlo, pCsBchlo, pCsCchlo and pCsEchlo) (Figure S9C&D). These vectors are reduced derivatives of pCk and pCs respectively, generated after removal of LB, RB and elements necessary for stability in Agrobacterium. For most chloroplast applications, L2 assemblies are sufficient. A L2 device is integrated into the chloroplast genome via homologous recombination. For customized targeted integration (Figure 3), positions L1-Ck1 and L1-Ck4 contain the homologous sequences of the native chloroplast genome, positioned upstream and downstream to the integration site. Positions L1-Ck2 and L1-Ck3 contain the transcription units for integration into the chloroplast genome.

**Figure 3.**
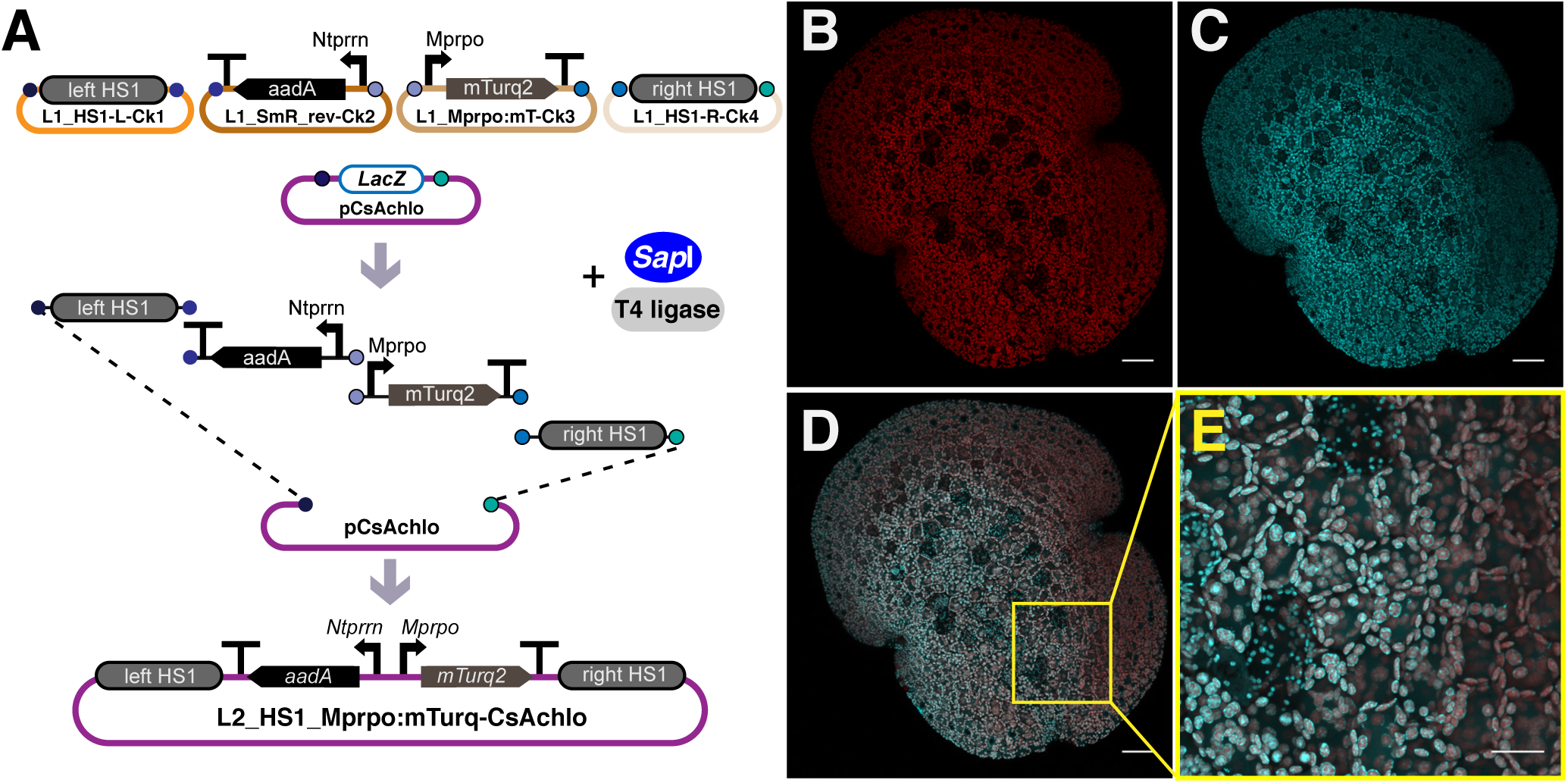
OpenPlant DNA tools for chloroplast genome modification. (A) Schematic representation of Level 2 Loop assembly of a device to express the chloroplast codon optimized mTurquoise2 fluorescent protein (mTurquoise2_chlo) under the control of the native Mprpo promoter using the homologous sequences left-HS1 and right-HS1 for integration in the chloroplast *trnG–trnfM* intergenic region. (B-E) Maximum intensity Z projection confocal images of Marchantia transplastomic gemmae transformed with the construct in A. (B) Chlorophyll autofluorescence channel, (C) mTurquoise2_chlo channel, and (D) merged channels. (E) Higher magnification images showing mTurquoise2_chlo accumulation inside the chloroplasts of all cells. Scale bar: 50 µm (B-D), 20 µm (E).

The OpenPlant kit contains left and right homologous sequences to allow integration in two intergenic regions (Figure S10). For integration in the *trnG*–*trnfM* region: the left homologous sequence (HS1-L, genome position: 40216 - 41843, Genbank accession no. MH635409) and right homologous sequence (HS1-R, genome position: 41847 - 43622, Genbank accession no. MH635409). For integration in the *rbcL*-*trnR* region: the left homologous sequence (HS2-L, genome position: 56139 - 57662, Genbank accession no. MH635409) and the right homologous sequence (HS2-R, genome position: 57681- 59304, Genbank accession no. MH635409). It should be noted that the homologous chloroplast sequences are derived from the Marchantia Cam-1/2 isolates, and the HS1 and HS2 sequences contain several nucleotide polymorphisms across their 1.5 kb lengths, compared to the Tak-1 isolate, for example.

The OpenPlant toolkit also contains the chloroplast codon optimized mTurquoise2 fluorescent protein coding sequence (mTurquoise2_chlo) ^22^ (Figure 3 and S10). In addition to the previously used promoters from *Nicotiana tabacum:* the photosystem II protein D1 gene promoter (pNt*psbA*) and the plastid ribosomal RNA operon promoter (pNt*prrn*) ^22^, the kit also contains the Marchantia native chloroplast DNA-dependent RNA polymerase operon promoter (pMp*rpo*) (genome position: 5591-5739, Genbank accession no. MH635409) (Figure 8). All parts have been functionally validated (Figure S10).

### DNA tools for nuclear genome editing

CRISPR-Cas9 systems can be used to introduce double strand breaks at target DNA sequences *in vivo* ^26, 27^. This allows the introduction of small deletion or insertion mutations via non-homologous end joining reactions at specific chromosome locations. CRISPR-Cas9-based targeted mutagenesis has been demonstrated in Marchantia ^28^, albeit with low frequency. This made the isolation of successful mutation events challenging. The efficiency of gene editing in plants has been improved by employing *Cas9* genes with altered codon usage ^29^. We chose a codon optimized Cas9 gene from the Puchta lab that was developed for use in *Arabidopsis thaliana* ^30^ and this has proved to have higher genome editing efficiency than an earlier version ^31^.

The OpenPlant kit contains a L2 vector (L2_lacZgRNA-Cas9-CsA) that allows Agrobacterium-mediated co-transformation of the Cas9 protein (expressed under the control of the p5-MpEF1α promoter) and a guide RNA (gRNA) expression cassette (under the control of the Marchantia U6 promoter) (Figure 4A). To target a gene, the corresponding gRNA sequence is directly cloned into the L2 vector using *Sap*I mediated Loop assembly (Figure S11A and Material and Methods). Dual gRNA editing can be achieved by combining into a L2, two L1 vectors with a gRNA expression cassette in each of them (Figure 4B-C). A gRNA target sequence is cloned into the L1 vector using *Bbs*I mediated Loop assembly (Figure 4B, S11B and Material and Methods). Dual gRNA editing allows, for instance, the generation of knockout mutants with longer DNA deletions by targeting two nearby sites in the genome (Figure S12).

**Figure 4.**
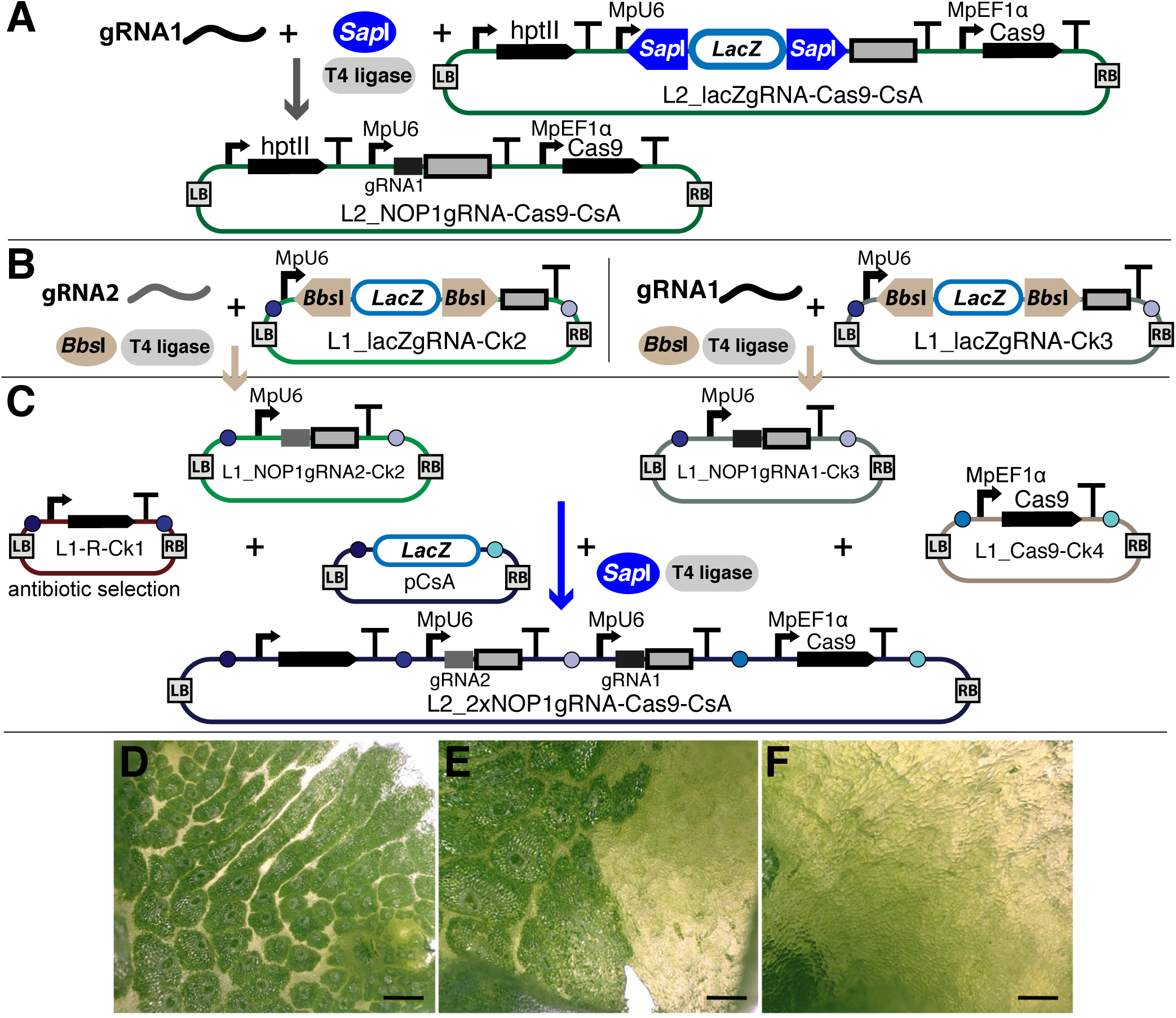
OpenPlant toolkit vectors for CRISPR/Cas9 genome editing. (A) Schematic representation of the plant transformation vector for simultaneous delivery of a single gRNA and Cas9 into Marchantia plants by Agrobacterium-mediated transformation. The gRNA sequence (in the example NOP1-gRNA1) was directly ligated into the L2_lacZgRNA-Cas9-CsA vector via *Sap*I type IIS assembly. (B-C) Schematic representation of assembly of the plant transformation vector for delivery of two gRNAs and Cas9 into a Marchantia plant. (B) The two gRNA sequences were ligated into L1_lacZgRNA-Ck2 and L1_LacZgRNA-Ck3 vectors, respectively, via *Bbs*I type IIS assembly. In the example, NOP1-gRNA2 into Ck2 and NOP1-gRNA1 into Ck3. (C) Then the two L1_gRNA transcription units were combined with an antibiotic resistance transcription unit and a MpEF1α:Cas9 transcription unit via *Sap*I Loop assembly. (D-F) The three phenotypic classes of regenerating Marchantia plants transformed with a L2 vector with NOP1gRNA and Cas9: wild type (D), chimeric (E) and mutant (F). Scale bar: 400 µm.

To test the efficacy of the new vectors (expressing either a single gRNA or a pair of gRNAs), we targeted the E3 ubiquitin ligase gene *Nopperabo1* (Mp*NOP1,* Mapoly0008s0036) gene. Mutation of this gene results in loss of air chambers, an easily scored and non-lethal phenotype ^32^. The delivery of either single or double gRNAs resulted in around half of the primary transformants showing mutant phenotypes. We observed three categories of regenerating sporelings, wild type, chimeric and mutant plants respectively (Figure 4D-F). The percentage of mutant plants with no observable air chambers was between 16-23 % of the total (Figure 4F, Figure S12 and Table S2). The OpenPlant toolkit also contains L0 parts with the Cas9 protein as CDS and the MpU6 promoter ^28^, allowing its users to create various devices for genome editing based on need (Figure 8).

### DNA tools for nuclear genome transformation

The Loop pCk and pCs vectors allow assembly of constructs that combine different gene elements and can contain multiple transcription units. In each Loop reaction, four vectors of the same level need to be ligated together, and thus, if wanting to assemble less than four units, for the vector(s) without a unit, a spacer will be used instead (an arbitrary 200 base pair DNA sequence). Spacer sequences are provided in the form of four pCk and four pCs plasmids. An example of use is shown in Figure 5A with pCk2 and pCk4. Loop assembly plasmids contain four ordered sites for sequence insertion, and the LB is adjacent to the first position, used for insertion of pCk1 or pCsA. Ideally, nuclear transformation vectors are designed with genes for antibiotic resistance located adjacent to the LB (Figure 5A), as Agrobacterium-mediated T-DNA transfer is initiated from the RB ^33^, and presence of a selectable marker at the distal terminus can improve the rescue of full-length insertions after plant transformation.

**Figure 5.**
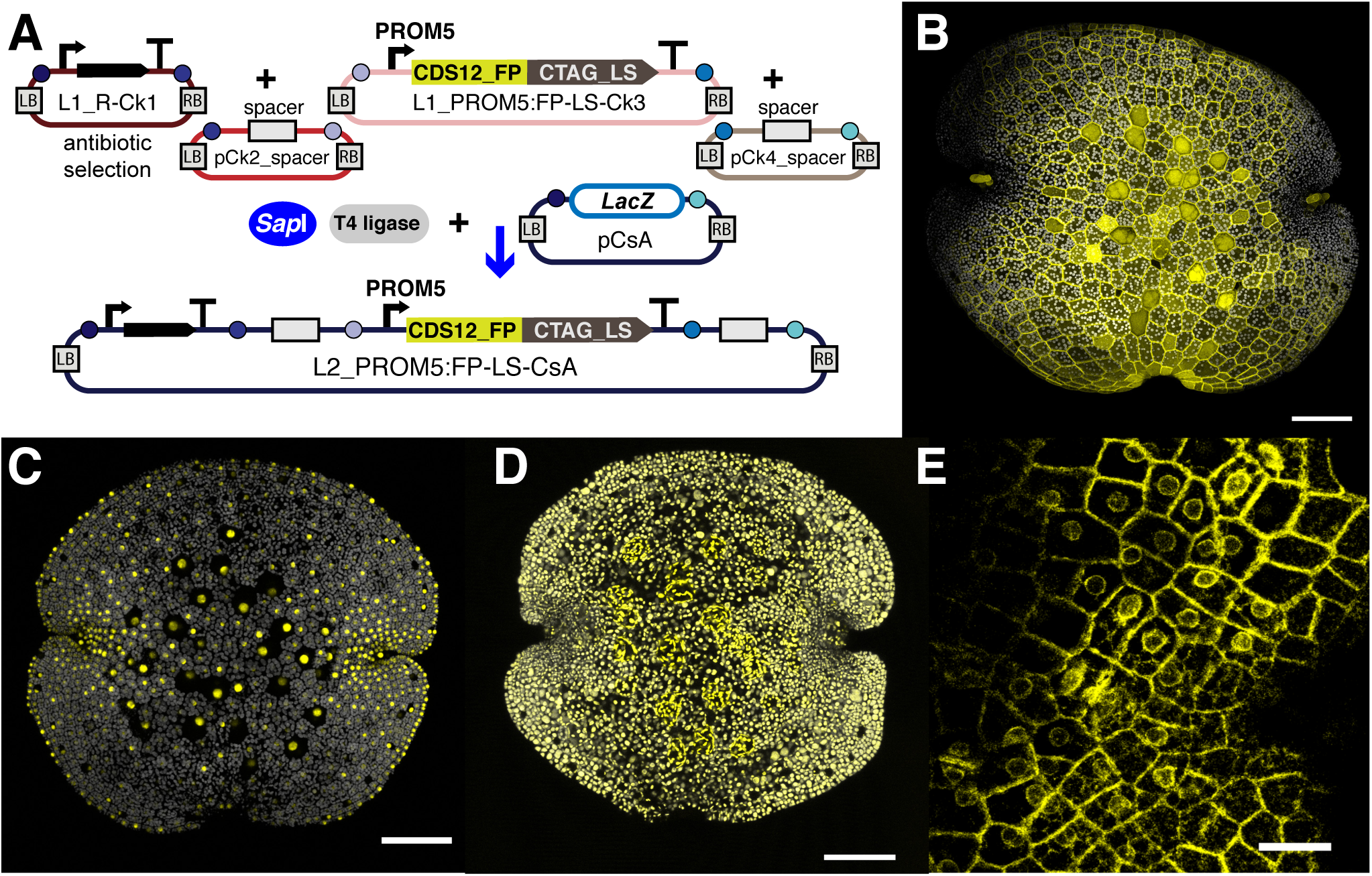
Testing different protein localization using the OpenPlant toolkit. (A) Schematic representation of the Level 2 Loop assembly of a construct for Agrobacterium-mediated nuclear transformation, built from Level 1 plasmids that contain an antibiotic resistance transcription unit in position 1 and gene for a fluorescent protein (CDS12_FP) fused to a localization signal (CTAG_LS) in position 3. Positions 2 and 4 contain spacer sequences. (B-E) Images of gemmae expressing different combinations of PROM5:FP-LS, with FP signal shown in yellow and chlorophyll autofluorescence in grey for all images. (B) p5-35S:mScarletI-LTI6b for plasma membrane localization (LTI6b is from the Arabidopsis gene AT3G05890). (C) p5-MpEF1α:mTurquoise2-N7 for nuclei localization (N7 is from the Arabidopsis ankyrin-like protein, AT4G19150). (D) p5-MpEF1α:MpSIG2-mVenus with chloroplast signal peptide from Marchantia SIG2 Mapoly0214s0004. (E) p5-35S:MpER-Targ-eGFP-HDEL with ER targeting peptide from a predicted Marchantia chitinase Mapoly0069s0092 (Figure S13). Images of gemmae at 0 d (B-D) or 4 d (E). Scale bar: 100 µm (A-C), 20 µm (D).

Reporter genes are essential components for constructing genetic circuits in new systems. Marchantia is phylogenetically distant from other model plants, and thus well tested components may need refactoring for use in this early divergent land plant. The OpenPlant toolkit contains L0 parts to assemble transcription units combining a wide range of multispectral fluorescent proteins (FPs) and targeting signals (Figure 5 and 8). We have systematically characterized and validated the performance of a series of modern fluorescent reporter genes. The kit contains genes for fluorescent proteins that are efficiently expressed and bright in Marchantia: eGFP ^20, 34^, mVenus ^16, 35^ mTurquoise2 ^16, 36^ and mScarlet-I ^37, 38^. mScarlet is a monomeric bright RFP developed from a synthetic template ^37^. Other monomeric RFPs, like tagRFP-T and mRuby3, have a residual tendency to dimerize, and carry additional problems due to incomplete or partial maturation ^37^. The fluorescent protein coding sequences are supplied as standardized parts (CDS12 or CTAG) to allow construction of N- and C-terminal protein fusions. We supply versions with or without a linker (SG10xAla) for translational fusion with proteins of interest, such as proteins that confer specific subcellular localization, or functional peptides. The kit includes five parts for protein fusion and targeting to different cellular compartments (Figure 5B-D). LTI6b from Arabidopsis ^39^ can be used to localize a protein to the plasma membrane ^16, 20^ (Figure 5B). Peptide sequences N7 from Arabidopsis ^16, 39^ and the simian virus 40 large T-antigen nuclear localization signal (SV40-NLS) ^20, 40^ confer nuclear localization (Figure 5C). The N-terminal fusion of the MpSIG2 signal peptide allows targeting of a fusion partner into the chloroplast^41^ (Figure 5D).

**Figure 6.**
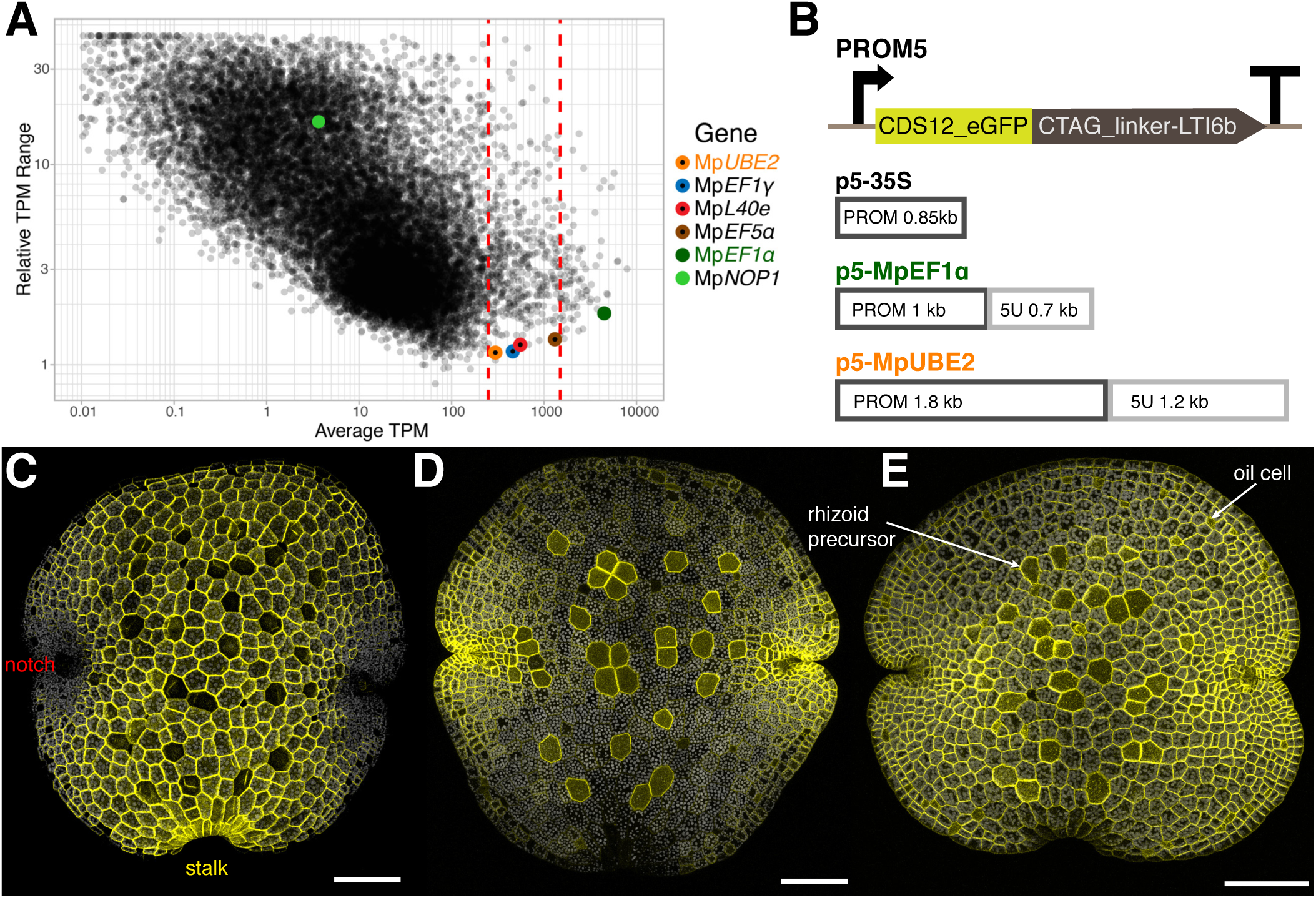
Identification of a ubiquitous and evenly expressed promoter for Marchantia gemmae. (A) Analysis of expression levels from publicly available RNAseq experiments for Mp genes. Variability of expression versus average expression levels for each gene (as relative TPM range [(maxTPM-minTPM)/avgTPM] versus average TPM). Genes with an average TPM in between 250-1500 are shown bracketed by dashed red lines. Colored dots represent 4 housekeeping genes that were tested as a source of constitutive promoters (including Mp*UBE2*). Dark and light green dots represent Mp*EF1α* and Mp*NOP1*, with average TPM of ∼4500 and ∼25 respectively. (B) Schema of transcription units for expression of plasma membrane localised eGFP. The eGFP-LTI6b gene was fused to the promoters: p5-35S (C) p5-MpEF1α (D) and p5-MpUBE2 (E). (C-E) Images of gemmae just removed from the gemma cup (0 days-old). Gemmae oriented with the two notch regions in the middle of the image and at the bottom, the region that was attached to the stalk in the gemma cup. There are two differentiated cell types on the surface of the gemmae that can be distinguished by their morphology: rhizoid precursors and oil cells (E). MpUBE2 is ubiquitously expressed at similar levels in all cells, while variable levels of expression were seen with the commonly used promoters 35S and MpEF1α. (C-E) eGFP is shown in yellow and chlorophyll autofluorescence in grey. Scale bar: 100 µm.

**Figure 7.**
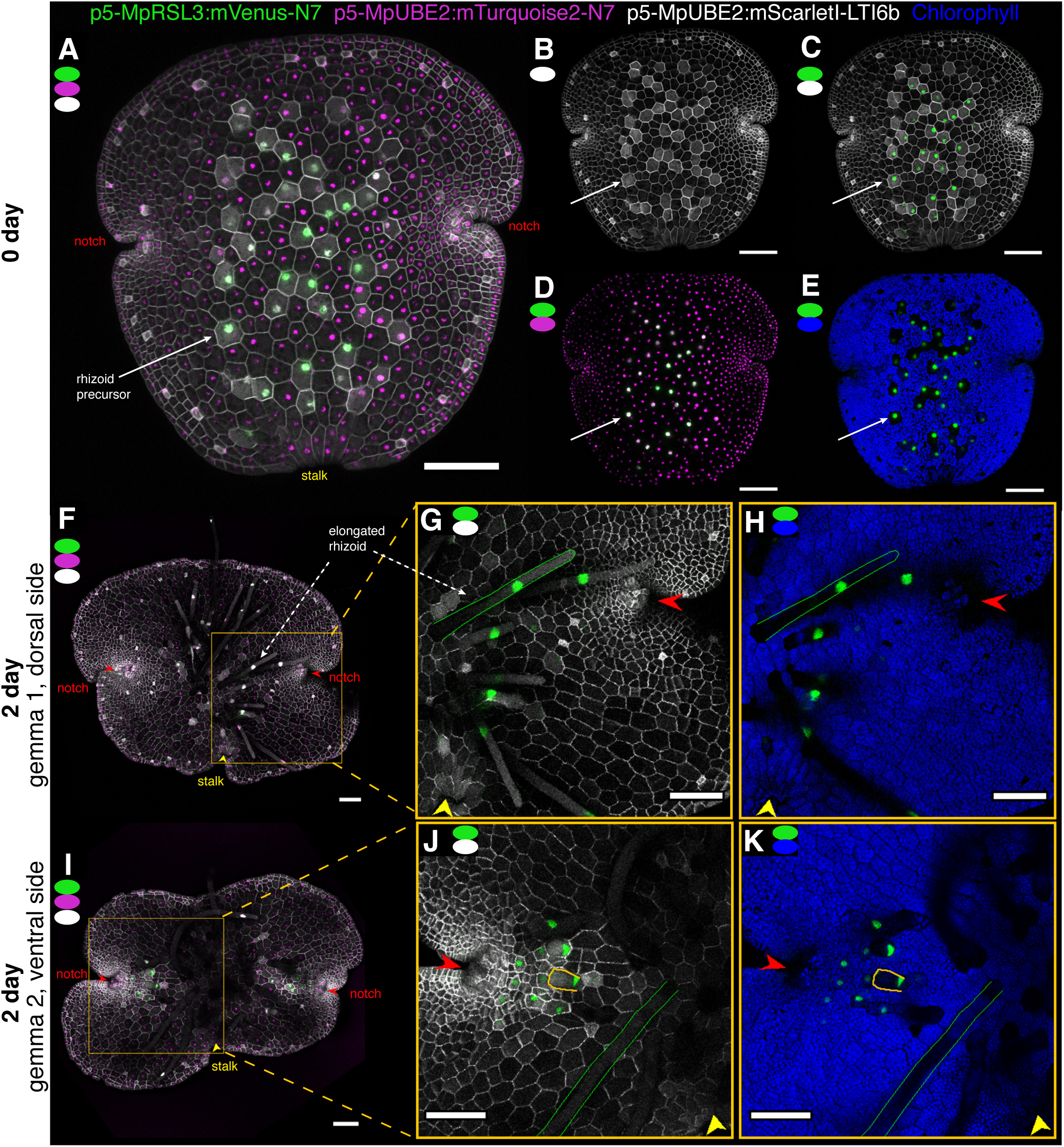
Construction of precise markers for visualizing cell differentiation using proximal promoter elements. The MpUBE2 and MpRSL3 proximal promoters confer ubiquitous and rhizoid specific expression, respectively, and were used to build gene assemblies for multispectral imaging of live gemmae. Maximum intensity z-axis projection images (A-E) of a gemma just removed from the gemmae cup (0 day). (A-E) Images from a line expressing a set of marker genes: p5-MpRSL3:mVenus-N7 (nuclear-localised rhizoid cell marker, green), p5-MpUBE2:mTurquoise2-N7 (nuclear-localised ubiquitous expression, magenta) and p5-MpUBE2:mScarletI-LTI6b (plasma membrane-localized ubiquitous expression, grey). (B-E) The relative levels and localization of gene expression can be dissected in separately combined image channels. (E) Chlorophyll autofluorescence, blue. (F-K) Images of whole 2 day old gemmae showing the dorsal (F) or the ventral side (I) (for details see Figure S14). Regions from the center of the gemmae to a meristem notch are shown enlarged (G,H,J,K). Rhizoid-specific expression (green) is shown with (grey) cell-outlines (G,J) and (blue) chlorophyll (H,K). The MpRSL3:mVenus-N7 marker is expressed in rhizoid precursor cells in the central part of the gemmae at day 0 (A-E), and in elongated rhizoids emerging from the central zone of the gemmae at day 2 (F-K). Hair-like rhizoids are outlined in green (G-H, J-K). Nuclei in elongated rhizoids move towards the tip of the cells (G-H). Expression of the MpRSL3 marker is triggered in cells near the notch, only on the ventral side of gemmae (J-K). These subsequently differentiate into rhizoid hairs (Figure S15). An early, bulged rhizoid cell outlined in orange. Scale bar: 100 µm.

**Figure 8.**
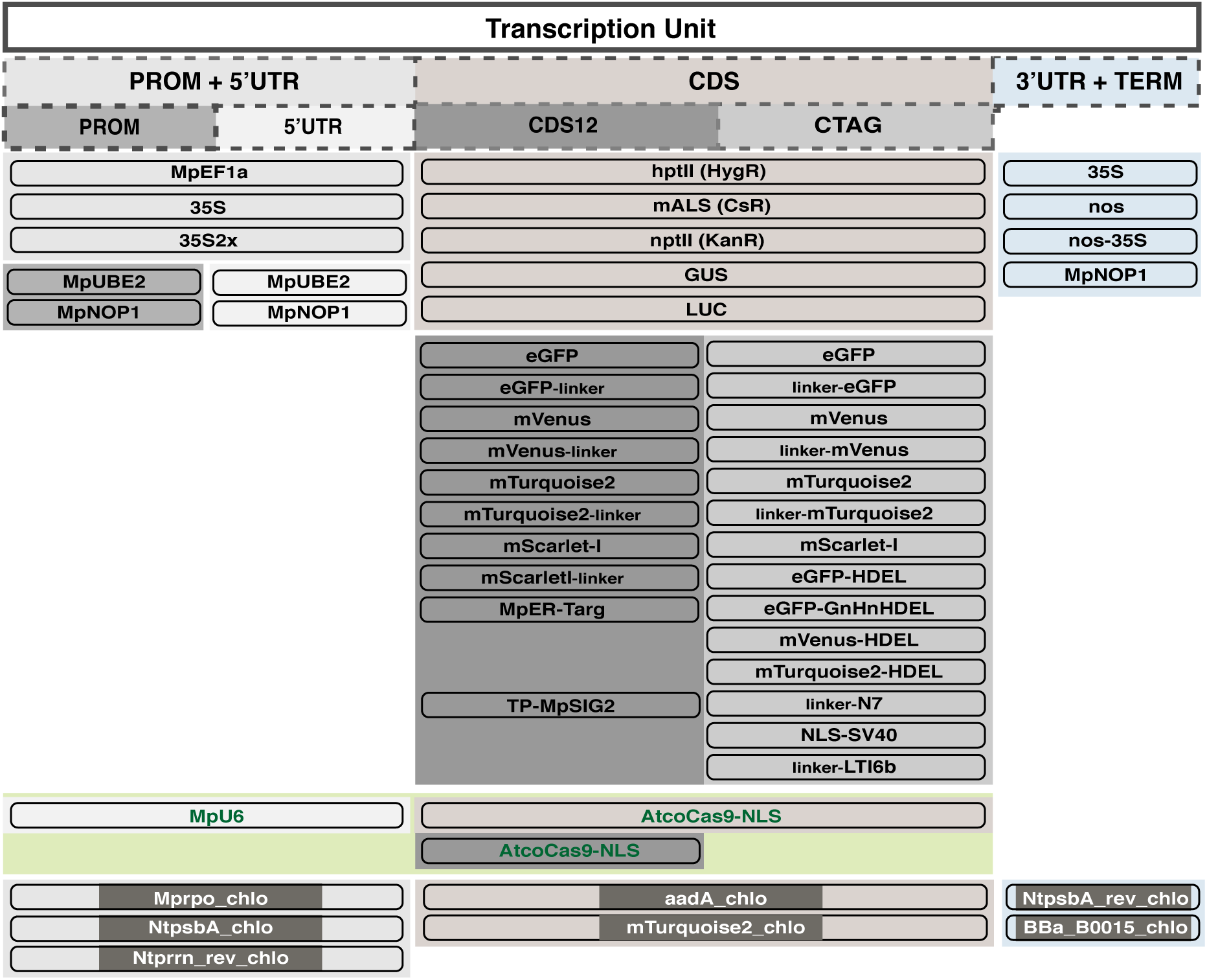
Level 0 DNA parts in the OpenPlant toolkit. The OpenPlant kit contains L0 parts to build transcription units, including: promoters with 5’ UTR (PROM5), promoters (PROM) and 5’ untranslated region (5UTR); coding sequences with start and stop codons (CDS), and for N-terminal (CTAG) and C-terminal (CDS12) protein fusions, and 3’ UTR with terminator (3TERM). The OpenPlant toolkit contains 50 L0 parts, 43 for nuclear transformation, 3 for CRISPR-Cas9 genome editing (in green), and 7 for chloroplast transformation (name_chlo).

The N-terminal signal peptide from Arabidopsis basic chitinase (AT3G12500), together with the C-terminal HDEL for protein retention in the ER, has been successfully used in different species of higher plants for ER localization ^42^. Nevertheless, when tested, it did not produce the expected pattern in Marchantia. We found three predicted native chitinases in Marchantia, tested the leader peptide from one of them, Mapoly0069s0092, fused to eGFP-HDEL and could observe fluorescence in reticulate structures in the cytosol, around the nuclei and in the plasma membrane (Figure 5E). In the OpenPlant toolkit we provide three different color FPs fused to HDEL as well as eGFP with a GlyHis-tag before HDEL as described before ^43^. The pumpkin 2S albumin leader peptide together with the ER retention signal HDEL has also been successfully used in Marchantia ^20^ and is compared with chitinase signal peptide sequences (Figure S13).

Besides fluorescent proteins, other systems are widely used to report promoter activity like β-glucuronidase (GUS) ^44^ and firefly luciferase (LUC) ^45^, and the OpenPlant kit contains as L0 CDS parts both the GUS and LUC genes (Figure 8).

### Synthetic promoters for nuclear genome transformation

Both for reference of constitutive expression and for extracting cell outlines, we need promoters expressed everywhere in the gemmae, at similar levels in all cell types, and with expression levels not too high, which could be toxic to the plant, or too low, which makes imaging more challenging.

Two constitutive promoters are commonly used in Marchantia, the cauliflower mosaic virus *35S* (CaMV *35S*) promoter and the endogenous *ELONGATION FACTOR 1*α (Mp*EF1α*, Mapoly0024s0116) promoter ^46^. There are two versions of the *35S* promoter that are used in Marchantia: p5-35S ^16, 47^ and p5-35Sx2 ^20^. Both of them have a strong activity in all parts of the gemmae and thallus except the notch area, where its activity is weak, and the oil body cells, where it is not active ^48^ (Figure 5B, 6C). On the other hand, the Mp*EF1*α promoter is expressed in all parts of the thallus, but it is especially strong at the notch area ^46^ (Figure 5C, 6D). Also, as shown in published transcriptome studies (Table S3) Mp*EF1*α expression levels are very high throughout development (Figure 6A and Table S3). We observed some slower and deformed growth and higher frequency of silencing events in primary transformants when using p5-MpEF1α to drive fluorescent protein expression. High levels of transcription driven by p5-MpEF1α seems to be not well tolerated by plants. We exploited our pipeline for efficiently building constructs, transforming sporelings and screening Marchantia gemmae, to test alternative promoters.

To identify new constitutive promoters without the limitations of the CaMV *35S* and Mp*EF1α* promoters, we reanalyzed publicly available Marchantia transcriptome data and screened for genes with medium to high levels of expression and minimal variability (Figure 6A, Table S3). As reference genes we used the strongly expressed Mp*EF1α* gene and the lowly expressed Mp*NOP1* gene ^32^. The average expression levels for Mp*EF1a* and Mp*NOP1* are around 4500 and 25 TPM (transcripts per million) respectively (Figure 6A, Table S3). We selected candidate genes with an average expression in between 250-1500 TPM and ranked them by their variability across all RNA seq experiments. We used the relative range as a measure of variability, calculated as (maximum TPM - minimum TPM) / average TPM. (Table S3). From the least variable genes, we selected four housekeeping genes with different average expression levels: the translation initiation factor 5a (Mp*EL5α*, Mapoly0045s0102, ∼ 1300 TPM), the large subunit ribosomal protein (*MpL40e*, Mapoly0003s0257, ∼ 560 TPM), the elongation factor 1γ (Mp*EF1*γ, Mapoly0062s0115, ∼ 460 TPM), and the ubiquitin-conjugating enzyme E2 (Mp*UBE2*, Mapoly0106s0019, ∼300 TPM) (Figure 6A and Table S2). Mp*EL5α* has been previously tested as a reference gene for qPCR ^49^. We extracted putative promoter sequences for those candidate genes: Mp*EL5α*, Mp*L40e*, Mp*EF1*γ all as a PROM5 part of 1.8 kb upstream ATG, and Mp*UBE2* as a PROM 1.8 kb part plus a 5U 1.2 kb part upstream ATG (Figure 6B). We assembled Loop L2 constructs with those promoters driving mTurquoise2 localized to the nuclei and tested them *in vivo*. Localization of the fluorescent gene product to the nuclei allowed easier comparison of levels of gene expression in different size cell types. p5-MpEL5α was weakly expressed in 0 day-old gemmae, only in the notch area and the rhizoid precursors, p5-MpL40e was expressed throughout the gemmae but with higher expression observed in nuclei in the region of the meristematic notch, and we could not detect expression driven by p5-MpEF1γ. p5-MpUBE2 was expressed in all cell types and throughout gemmae development both when driving mTurquoise2 to the nuclei and eGFP to the plasma membrane (Figure 6E). Thus, we selected MpUBE2 promoter to drive constitutive expression of transgenes in early stages of gemmae development.

We then tested the new constitutive promoter p5-MpUBE2 in multispectral imaging, for reference of expression and to ubiquitously label cell boundaries together with a marker for specific cell types. We made L2 devices containing transcription units composed of the p5-MpUBE2 driving the expression of mTurquoise2 localized to the nuclei, the p5-MpUBE2 driving the expression of mScarletI localized to the plasma membrane, as well the MpRSL3 proximal promoter (as a PROM5 1.8 kb part) driving the expression of mVenus targeted to the nuclei (Figure 7). Mp*RSL3* (MpbHLH33, Mapoly0029s0135) is a transcription factor (TF) from the subfamily VIIIc of basic helix-loop-helix TFs, which in Marchantia has three members ^50^. One of them, MpRSL1 (MpbHLH14, Mapoly0039s0003) has been identified as a ROOTHAIR DEFECTIVE SIX-LIKE (RSL) class I gene and it is required for the initiation and development of unicellular rhizoids, mucilage papillae and gemmae ^51, 52^. When imaging gemmae just displaced from the gemma cup (0 days-old gemma), MpRSL3 promoter expression could be observed in the rhizoid precursor cells (Figure 7A-E). Both sides of a 0 days-old gemma have rhizoid precursor cells ready to elongate as rhizoid cells when landing. Once gemmae are displaced from the gemmae cup ^6^, dormancy is released and dorsiventral polarity is specified with formation of air chambers on the dorsal side and formation of rhizoids on the ventral side (Figure 7 and S14). We tracked gemmae for 2 days, and the MpRSL3 promoter kept expressing after the rhizoid precursors bulged and became elongated rhizoids with their nuclei moving towards the tip of the cells (Figure 7F-K, Figure S15). On the dorsal side of a gemmae, the MpRSL3 marker was not expressed in new cells and no new rhizoid precursor cells were specified (Figure 7F-H, Figure S15). On the other hand, mTurquoise2 expressed under the control of the MpUBE2 promoter accumulated in all cells (Figure 7, S15). To image the ventral side of gemmae, we grew them in inverted plates (Figure S14). On the ventral side, we observed that new cells near the notch started expressing the MpRSL3 promoter and became rhizoid precursors which subsequently elongated (Figure 7I-K, S15). Thus, the MpRSL3 proximal promoter can be used to study the dynamics of rhizoid precursors specification and elongation (position, number, spacing, timing), in combination with markers for reference of expression and ubiquitous labeling of cell boundaries under the control of the MpUBE2 promoter. Constructs like the one used here will advance the understanding of dynamic changes in expression patterns in Marchantia, as a first step to screen for regulatory sequences to engineer morphogenesis in this simple plant system.

## CONCLUSIONS

The development of systematic approaches to the engineering of microbial genetic systems has facilitated substantial advances in science and biotechnology. There are many potential benefits in applying the same approach in plant systems. However, work with plants is beset with problems due to slow generation times, genetic redundancy and difficulties in experimental manipulation and quantitative observation, compared to microbes. We have set out to establish a new technical platform for facile engineering of plants, to circumvent many of these difficulties.

We have adopted *Marchantia polymorpha* as a simple plant system that has an open form of development, which allows direct visualization of tagged gene expression and cellular growth in living tissues. This liverwort plant system is fast and easy to culture and transform. Here, we describe methods to maintain the plant in axenic culture throughout its life cycle, using a combination of growth in agar-containing Petri dishes and on Jiffy-7 peat disks in Microboxes^TM^. We also describe methods for cryopreservation of Marchantia gemmae, to enable easier maintenance and storage of large numbers of plant lines. These streamlined methods allow routine propagation of Marchantia and provide a simple source of sterile propagules for manipulation or observation. These simple culture techniques allow laboratories to tackle advanced bioengineering projects in plants without the need for glasshouse facilities. We believe that the adoption of Marchantia as a prototype will widen access to synthetic biology experiments in plants.

Marchantia tissues show an extreme ability to regenerate without external cues. Regenerated plants spontaneously produce clonal propagules within a few weeks of an experiment. Patterns of gene regulation and cellular dynamics can be directly analyzed in these propagules. It has a simple cellular architecture and streamlined genome with highly reduced gene redundancy. The plant is haploid, and has a genome of around 220 Mb and 19,000 encoded genes^7^. Many families of regulatory genes are found with highly reduced numbers compared to higher plants, and whole tissue transcriptomic data is available for gene expression at different stages of growth. We have established a common genetic syntax for the composition (and exchange) of plant DNA parts (Phytobricks), recruiting wide support from the international plant science community ^15^. Further, MarpoDB was built as an open source database that handles the Marchantia genome as a collection of potential Phytobrick parts ^54^. This is highly useful for automatic mining of new parts, and managing part characterization.

We describe the modification of a published method for gene assembly ^15, 16^. Loop assembly is based on recursive use of two Type IIS restriction endonucleases, and allows rapid and efficient production of large DNA constructs, is compatible with widely used Level zero (L0) DNA parts such as Phytobricks, and can be easily automated. We have constructed an integrated set of plant transformation vectors for transformation and editing of the nuclear genome, and modification of the chloroplast genome. The new vectors are easier to handle in the laboratory, and provide new features. The vectors maintain compatibility with modular DNA parts that follow the Golden Gate, MoClo and Phytobrick syntax, and we have built and tested a wide range of new parts. These include a wide range of gene cassettes for multispectral fluorescent proteins, gene control, Cas9 and sgRNA expression, plastid gene regulation and sequences for site-specific insertion into the Marchantia chloroplast genome by homologous recombination. The vectors and part collections are being made available through the Open Materials Transfer Agreement (OpenMTA ^17^). Use of the OpenMTA places the materials into the public domain, allowing redistribution of materials and use for commercial projects.

Marchantia gemmae provide a new biological system with unique advantages for genetic reprogramming of plant growth. Marchantia plants produce gemmae spontaneously, as vegetative clonal propagules. New gemmae are formed around 2-3 weeks after germination of a vegetative parent gemma. New gemmae form inside conical splash cups and can be harvested prolifically. Marchantia gemmae have a stereotypical and simple cellular architecture. They can be germinated and grown directly under microscopic observation. Gene expression can be visualized at the scale of individual cells. Dynamic cellular relationships within growing tissues can be directly visualized and quantified. Both short-range and organism-wide coordination of growth and differentiation can be perturbed and mapped in a way that is difficult or impossible in other plant systems due to tissue complexity, optical inaccessibility, genetic redundancy or all of the above. We have employed the new toolset described here to build and test new markers for visualizing cell dynamics.

An important aspect when engineering multicellular organisms, is the ability to target expression to specific cell types, at specific times in development, and at the desired expression levels. We have demonstrated the use of genome annotation and transcriptome data to design candidate promoter elements, and successfully characterised a new promoter MpUBE2, that drives ubiquitous gene expression in gemmae. Transcription factors initiate and maintain the transcriptional programs that specify the diversity of cell types in an organism and control growth. Marchantia has a low genetic redundancy, all main plant transcription factor families are comprised of only ∼400 genes. We are conducting a systematic screen of TF promoter fusions in Marchantia using the tools for high-throughput transformation and fluorescence microscopy described here (Figure 1D, Figure S13, materials and methods). We believe that these new methods and the compound benefits of Marchantia as an experimental system will allow new approaches to important and fundamental questions about plant growth and engineering.

We have developed the OpenPlant toolkit as a set of interlinked resources and techniques, which have been designed to facilitate and standardize DNA assembly, and manipulation of both nuclear and chloroplast genomes, thus enabling the engineering of Marchantia development and metabolism. We hope that the tools developed in this work will encourage the adoption of Marchantia as a simple multicellular prototype for bioengineering, lower the logistical threshold for plant experiments and promote the recruitment of non-botanists to plant synthetic biology research, as well as encourage open science approaches to support standardization, reproducibility and innovation.

## MATERIALS AND METHODS

### Plant material and growth conditions

*Marchantia polymorpha* accessions Cam-1 (male) and Cam-2 (female) were used in this study ^54^. Plants were grown and maintained on 0.5x Gamborg media (0.5x strength Gamborg B5 medium plus vitamins, Duchefa Biochemie G0210, pH 5.8) and 1.2% (w/v) agar (Melford capsules, A20021), under continuous light at 21 °C with light intensity of 150 µmol/m^2^/s.

### Maintenance and storage of lines

To maintain a line, Marchantia plants can be grown under continuous light at 14 °C with light intensity below 100 µmol/m^2^/s. In this way plants can be kept for two-three months before transfer to new plates. Gemmae can be stored from 6 months to a year at 4 °C in the dark in 1.5 mL Eppendorf tubes half filled with 0.5x Gamborg media 1% (w/v) agar.

For longer term storage, lines can be cryopreserved at −80 °C for at least 1-2 years^18^. We simplified the published cryopreservation protocol for *Marchantia polymorpha* gemmae^18^. Gemma were placed on preculture plates and incubated for 1-3 days under normal growth conditions. Preculture plates consisted of 0.5x Gamborg media, 1.2% (w/v) agar, 0.3 M sucrose, 10 µM ABA. Media was autoclaved without ABA, and ABA added from a filter-sterilized stock solution just before pouring plates. For encapsulation of gemmae, small drops (approx. 40 µL) of alginate solution (0.5x Gamborg media, 3% sodium alginate) were placed on an empty 4.5 cm petri dish to form beads. Gemmae were transferred from preculture plates to the petri dish using forceps; up to five gemmae were placed inside each bead. A drop of CaCl2 solution (0.5x Gamborg B5 media, 0.1 M CaCl2) was added, and beads were allowed to solidify for 10 min. The plate was filled by serological pipette with dehydration buffer (0.5x Gamborg B5 media, 2 M glycerol, 1 M sucrose), submerging the beads. Beads were soaked for 30 min, and afterwards the dehydration buffer was removed with a serological pipette without disrupting the beads. The lid was removed and the beads were air dried in the flow hood for at least 2 hours. Using forceps or scalpels the beads were transferred into 1.5 mL Eppendorf tubes (max. 10 beads/ tube). Tubes were flash frozen in liquid nitrogen for 2 min and stored at −80 °C. For thawing and recovery of lines, tubes were removed from the freezer and immediately placed in a 37 °C water bath or heat block for 2 min. 1.5 mL of thawing solution (0.5x Gamborg B5 media, 1 M sucrose) was added to each tube, and the tubes were incubated at room temperature for 10 min. Then the thawing solution was replaced with rinse solution (0.5x Gamborg B5 media, 0.03 M sucrose) and the tubes were incubated at room temperature for 10 minutes. Beads with gemmae were placed on 0.5x Gamborg B5 plates, and incubated under normal growth conditions (Figure S4).

### Marchantia cultures for spore production

For spore production, 15 Jiffy-7 dehydrated peat disks were placed in a TP5000+TPD5000 Microbox^TM^ micropropagation containers with two #40 green filters (SacO2, Belgium), 800 mL of water added and the container autoclaved. After autoclaving, thallus fragments or gemmae were placed on each hydrated Jiffy-7 pellet, and another 200 mL of water added. Microboxes were placed at 21 °C, continuous light, 150 µE. After one month, Microboxes were transferred to long day conditions (16 hours light / 8 hours dark), with light intensity of 150 µE supplemented with far-red light (peak emission around 730-740 nm, from Philips GreenPower LED HF far red, or Intelligent LED Solutions ILS-OW06-FRED-SD111 730 nm). After approximately 4 weeks mature male and female reproductive organs developed. For fertilization, one drop of water was transferred, with a disposable sterile plastic pipette, from the male reproductive organs on the top of the female reproductive organs. After one more month mature sporangia were visible and ready to be collected in large quantities and stored with desiccant at 4 °C for 1-2 months or at −80 °C for longer storage (Figure S1).

### Sterilization of Marchantia spores

Sterilization solution was prepared as previously described ^7^, by dissolving one Milton mini-sterilizing tablet (Milton Pharmaceutical UK, Cheltenham, UK) in 25 ml of sterile water, so the final concentration of the active ingredient, sodium dichloroisocyanurate (NaDCC) was 0.05% (w/v). Sporangia were transferred to a 1.5 mL Eppendorf tube. For Agrobacterium-mediated transformation with multi-well microplates, two sporangia were used per well-transformation (e.g. 12 sporangia for 6 transformations in a 6-well plate). Sterilization procedure was as follows. Sporangia were transferred to a 1.5 mL Eppendorf tube and metal tweezers were used to crush them. 0.5 mL of sterilization solution was added to each tube and tubes vortexed to get a finely pulped suspension. A 40µM cell strainer was placed over a 50 mL falcon tube, and the 0.5 mL of crushed sporangia poured onto the filter. The eppendorf tube was rinsed with 1 mL of fresh Milton solution and poured onto the filter. The filter was washed with 2.5 mL of fresh Milton solution. The filtrate containing the spores (total volume approx 4 mL) was divided between four new 1.5 mL eppendorf tubes. Spores were sterilised in the sterilization solution for 20-30 min. Tubes were then centrifuged at 16000g for 2 min, the supernatant discarded and spores pooled into a single tube, re-suspending the pellet in 25 µL for each planned transformation. Spores were plated on 0.5 x Gamborg plates using 50 µL per plate (each plate provides enough material for 2 transformations). Plates were sealed with micropore tape and incubated (in an inverted orientation to prevent rhizoid growth into the agar media) for 5-7 days at 21°C under continuous light.

### Agrobacterium-mediated transformation in multiwell dishes

A modification of the published Agrobacterium-mediated sporeling transformation protocol ^19^ was used. See Figure S2 and protocols.io/OpenPlant. For each construct of interest, a single colony of Agrobacterium, transformed with the plasmid of interest, was inoculated into 5 mL LB media plus the antibiotics required for selection, and incubated at 28°C with shaking at 150 rpm for two days. The Agrobacterium cultures were centrifuged for 10 min at 2000 xg, resuspended in 5 mL of 0.5x Gamborg B5, 1% (w/v) sucrose, 100 µM acetosyringone (3′,5′-dimethoxy-4′-hydroxyacetophenone, Sigma D134406), and incubated for 6h at 28°C at 150 rpm.

In each well of a 6-well plate, Marchantia 5-7 day old sporelings were co-cultivated with 100 µL of Agrobacterium culture in 4 mL of liquid 0.5x Gamborg B5 plus supplements (0.1% N-Z amino A (Sigma C7290), 0.03% L-glutamine, 2% (w/v) sucrose) 100 µM acetosyringone media. The 6-well plate was then placed on a shaker at 120 rpm for 2 days at 21 °C with continuous lighting (150 µmol/m^2^/s). For each well, the sporelings were washed with 50 mL of sterile water and plated on 0.5x Gamborg B5 media plus antibiotics plates. Plates contain 100µg/ml cefotaxime and an antibiotic for selection of the transformant of interest; hygromycin 20 µg/mL (Invitrogen 10687010), chlorsulfuron (10 µM, Sigma 64902-72-3) or kanamycin (125 µg/mL, Sigma 25389-94-0).

### Preparation of Seashell nanoparticles for chloroplast transformation

Before bombardment, macro-carriers with the macro-carrier holders were placed in a glass Petri dish. Rupture disks and stopping screens were placed in another glass Petri dish. Both dishes were wrapped with aluminium foil and autoclaved. DNAdel^TM^ nanoparticles (Seashell Technologies) were supplied as a 50 mg/mL suspension in binding buffer. To dissociate any aggregates prior to use, the suspension was agitated, sonicated briefly for 30 s using an ultrasonic water-bath sonicator and vortexed for 5 s twice. 0.5 mg of nanoparticles per shot (transformation) were used. DNAdel^TM^ nanoparticles were diluted into Binding Buffer (Seashell Technologies) to final concentration 30 mg/mL in a 1.5 mL Eppendorf tube. 1.5-2 µg of plasmid DNA per shot was added into the tube. Finally, an equal volume of Precipitation Buffer (Seashell Technologies) was added into the tube (volume = volume of DNAdel^TM^ nanoparticles + volume of Binding Buffer + volume of plasmid DNA). The tube was vortexed and the mix was incubated at room temperature for 3 min. The tube was centrifuged at 8000 xg for 10 s, the supernatant discarded, and the DNA coated DNAdel^TM^ nanoparticles were washed with 500 µL ice cold 100% EtOH. The tube was centrifuged at 8000 xg for 10 s again, the supernatant discarded, and the nanoparticles were re-suspended in 100% EtOH with a volume of n times 7 µL, n= number of shots planned. To resuspend, the nanoparticles were briefly sonicated using an ultrasonic water-bath sonicator, usually two rounds of 5 s sonication. 7 µL of DNA coated nanoparticles was pipetted into the center of each macro-carrier and left to dry. Biolistic bombardment was carried out with the Bio-Rad PDS-1000/He device according to manufacturer’s directions. The tissues used for bombardment were sporelings grown for 7 days on inverted 0.5x Gamborg media plates. After bombardment, the plates were cultured in a growth room for 2 days, and then transferred to 0.5x Gamborg B5 plates with spectinomycin (500 µg/mL). After 4-6 weeks successful transformants were visible (Figure S3).

### Chloroplast genome

Illumina short reads of Cam-1 nuclear, mitochondrial, and plastid DNA were sorted against a Tak-1 draft assembly generously provided by John Bowman, and reads that matched the chloroplast genome were assembled *de novo* using CLC Genomics workbench. *De novo* assembly generated 25 contigs spanning the entire plastome, which were assembled using the reference Tak draft assembly to generate the complete plastome. The assembly process was then repeated with Illumina reads from Cam-2, and the assembled sequences were identical. The Cam-1/2 sequence has been submitted to GenBank with accession number MH635409.

### Genotyping

For genotyping ^22^, small pieces (3×3 mm) of thalli from individual plants were placed in 1.5 mL Eppendorf tubes and crushed with an autoclaved micro-pestle in 100 µl genotyping buffer (100 mM Tris-HCl, 1M KCl, 1M KCl, and 10 mM EDTA, pH 9.5). The tubes were then placed at 80 °C for 5 min. 400 µL of sterile water was added to each tube. 5 µL aliquots of the extract were used as a template for PCR and products were checked on a 1.5% (w/v) agarose gel.

### L0 reaction protocol

To clone a L0 part into the pUAP4 vector, the following steps were followed. Primers were designed as described in Figure S5A. DNA parts were PCR amplified from the source DNA part (e.g. plasmid DNA or genomic DNA), using a high fidelity DNA polymerase. PCR products were run on a 1.5% (w/v) agarose gel and the QIAquick Gel Extraction Kit used to extract the band that corresponds to the size of the amplified DNA part. Aliquots of the DNA part were prepared at a concentration of 15 nM and of the pUAP4 acceptor vector at a concentration of 7.5 nM. A type IIS assembly reaction was set up into a 0.2 mL tube (5 µL nuclease-free H2O, 1 µL 10x Tango Buffer (Thermo Fisher), 0.5 µL mg/mL bovine serum albumin (NEB), 0.25 µL T4 DNA Ligase at 5 U/ µL (Thermo Fisher), 1 µL 10 mM ATP (Sigma Aldrich), 0.25 µL *Sap*I (*Lgu*I) at 5 U/µL (Thermo Fisher), 1 µL of L0 DNA part and 1 µL of pUAP4) to clone the amplified DNA part into pUAP4. Samples were incubated in a thermocycler using the following program. Assembly: 26 cycles of 37 °C for 3 min and 16 °C for 4 min. Termination and enzyme denaturation: 50 °C for 5 min and 80 °C for 10 min. 20 µL of chemically competent *E. coli* cells (transformation efficiency of 1 × 10^7^ transformants/µg plasmid DNA) were transformed using 2 µL of the assembly reaction and then spread on LB agar plates containing 25 µg/mL chloramphenicol and 40 µg/mL X-gal. The presence of the correct insert was confirmed with Sanger sequencing using the primers UAP_F and UAP_R and any additional DNA part specific primers (Figure S5).

### L1 *Bsa*I mediated Loop assembly

L1 reactions were performed as previously described ^16^ with minor modifications. Briefly, L0 plasmids with DNA parts to be assembled were prepared at a concentration of 15 nM and the acceptor pCk vector at a concentration of 7.5 nM. Loop assembly Level 1 reaction master mix (MM) contained: 3 µL nuclease-free H2O, 1 µL 10x T4 DNA ligase buffer (New England Biolabs (NEB)), 0.5 µL 1 mg/mL bovine serum albumin (NEB), 0.25 µL T4 DNA ligase at 400 U/µL (NEB), 0.25 µL *Bsa*I at 10 U/µL (NEB). Cycling conditions were: 26 cycles of 37 °C for 3 min and 16 °C for 4 min. Termination and enzyme denaturation: 50 °C for 5 min and 80 °C for 10 min. 20 µL of chemically competent *E. coli* cells were transformed using 2 µL of the Loop assembly reaction and plated on LB agar plates containing 50 µg/mL kanamycin and 40 µg/mL of X-Gal. The presence of the correct insert was confirmed with Sanger sequencing using primers pC_F and pC_R for pCk vectors and pC_F and pC_R2 for pCkchlo vectors (Figure S7).

### L2 *Sap*I mediated Loop assembly

The protocol is the same as the L1 *Bsa*I assembly protocol with the exception of MM composition: 2 µL nuclease-free H2O, 1 µL 10x Tango Buffer (Thermo Fisher), 0.5 µL mg/ mL bovine serum albumin (NEB), 0.25 µL T4 DNA ligase at 5 U/µL (Thermo Fisher), 1 µL 10 mM ATP (Sigma Aldrich), 0.25 µL *Sap*I (LguI) at 5 U/µL (Thermo Fisher) and plated on LB agar plates containing 100 µg/mL spectinomycin and 40 µg/mL of X-Gal. The donor plasmids were L1 constructs and the acceptor plasmid was a pCs vector. The presence of the correct insert was confirmed with Sanger sequencing using the primers pC_F and pC_R for pCs vectors and pC_F and pC_R2 for pCschlo vectors.

### Loop automation using Labcyte Echo 550

L1 and L2 assembly reactions were automated and miniaturized for assembly of 500 nL reactions using the Echo 550 acoustic-focusing non-contact liquid handling robot from Labcyte (San Jose, CA, USA). For L1 reactions an LDV source plate (Labcyte) with all necessary L0 plasmids was needed. For L2 plasmids another LDV plate with all L1 plasmids was provides. We used a PP source plate (Labcyte) for acceptor plasmids (AP) as well as master mix (MM) and water. Calibrations 384LDV_AQ_B2 and 384PP_AQ_BP2 were used, respectively. Both LDV and PP plates were sealed and stored at −20°C for future use.

The assembly reaction conditions were identical to those described above for a final volume of 10 µL, but scaled down to a total volume of 500 nL. Plasmid DNAs in the source plates were 10x more concentrated than in the final reaction volume, and thus for each plasmid in a reaction/well the volume of plasmid to transfer to was 50 nL. For L1 reactions we used up to 6 L0 plasmids, the volume of plasmids was 300 nL and the volume of L1-MM was 200 nL (with these components in each well: water: 100 nL, 10x T4 DNA ligase buffer: 50 nL, 1 mg / mL bovine serum albumin (NEB): 25 nL, T4 DNA ligase at 400 U / µL (NEB): 12.5 nL, *Bsa*I at 10 U / µL (NEB): 12.5 nL). For L1 reactions assembling less than 6 plasmids, the volume was adjusted with water. For example, the volume of plasmid DNA in an L1 assembly with 4 plasmids (1 AP and 3 L0 plasmids) was 200 nL, and 200nL L1-MM and 100 nL water were added to the final reaction. All L2 reactions involved assembly of 5 plasmids (1 AP and 4 L1 plasmids) with a total volume of 250 nL, added to 250 nL L2-MM (100 nL water, 10x Tango Buffer (Thermo Fisher): 50 nL, 1 mg/mL bovine serum albumin (NEB): 25 nL, 10 mM ATP (Sigma Aldrich): 50 nL, T4 DNA ligase at 5 U/µL (Thermo Fisher) : 12.5 nL, *Sap*I (*Lgu*I) at 5 U/µL (Thermo Fisher): 12.5 nl) in each well. We first prepared the final volume of MM needed ((volume/well x number of wells) plus dead volume of PP wells) in an Eppendorf tube, mixed by pipetting and then transferred the MM to a PP-well. We created csv files in the Labcyte Cherry Pick software to add all the necessary information (source and destination wells for each sample and reaction, calibration and volume).

Before running the Loop reaction, source plates were defrosted, spun down and warmed at room temperature (∼22 °C). MM was prepared fresh, added to the PP plate, spun down and kept at room temperature. Reactions were set up in 384 well PCR plates (4titute 4ti-0384), sealed with foil (4titude 4ti-0550), spun down and incubated on a 384 well thermal cycling instrument using the conditions described above, with a lid temperature of 70 °C. Reactions were transformed into 2 µL of Top10 chemically competent cells (transformation efficiency of 1 × 10^8^ transformants / µg plasmid DNA). Heat shock was performed as follows: 2 µL of cells were added to each well, the plate was incubated 15 m on ice, incubated at 42 °C for 40 s in a thermocycler, and cooled for 2 min on ice. 15 µL of LB media was added to each well, the plate was sealed with a breath-easy sealing membrane (Z30059-1PAK) and incubated at 37 °C for 1 hour. Then the whole volume in each well was plated in selective LB agar plates as described above.

### Guide RNA cloning for gene-editing vectors

Two oligos for cloning of the gRNA target sequence were designed as described in Figure S11. 1 µL of each oligo (100µM) were mixed with 8 µL water into final volume of 10 µL and then annealed in a thermocycler with the following parameters: 37 °C for 30 min, 95 °C for 5 min and then ramped down to 25 °C at 5 °C per min. After annealing, the gRNA target sequence was cloned into the L2_lacZgRNA-Cas9-CsA vector using L2 *Sap*I mediated Loop assembly (Figure S11A). For cloning into the L1_lacZgRNA-Ck2 or L1_lacZgRNA-Ck3 vectors (Figure S11B), a *BbsI* mediated type IIS assembly was used: 6 µL nuclease-free H2O, 1 µL 10x T4 ligase buffer (NEB), 0.5 µL mg/mL bovine serum albumin (NEB), 0.25 µL T4 DNA ligase at 400 U/µL (NEB), 0.25 µL *Bbs*I (*Bpi*I) at 5 U/µL (Thermo Fisher), 1 µL vector and 1 µL annealed oligos.

### Analysis of public RNA-sequencing data

Fastq files for the following BioProject study accessions: PRJDB4420 ^55^, PRJDB5890 ^56^, PRJDB6579 ^57^, PRJDB7023 ^7^, PRJNA218052 ^58^, PRJNA265205 ^59^, PRJNA350270 ^7^, PRJNA397394 ^60^, PRJNA433456 ^61^, PRJNA251267 ^7^ were downloaded from the European Nucleotide Archive. FastQC was used to inspect read quality and TrimGalore to trim low-quality reads and remaining sequencing adapters. Reads were pseudo-aligned using kallisto ^62^ to the Marchantia Genome ^7^ version JGI3.1 (primary transcripts only, obtained from MarpolBase). Kallisto estimates abundance of each transcript in units of transcripts per million (TPM). We merged replicates by averaging and generated a gene-sample matrix of TPM values (Table S2). This matrix was used to calculate the average expression and relative range for each gene.

### Screening of transformants with fluorescent microscopy

Primary transformants growing on 0.5x Gamborg plates with antibiotic were screened using a stereo microscope with fluorescence (Leica M205 FA). Ten days after transformation and 20 plants were transferred onto new selection plates. We performed a second screen for fluorescence at the stereo microscope, selecting around 10 plants with expected signals. In about 2 more weeks, the plants produced gemmae cups containing G1 (generation 1) gemma. For each construct, we selected at least three independent lines to image gemmae with laser scanning microscopy using either slides or 384 well plates.

The Leica M205FA was equipped with the following filters for observation of fluorescence: ET CFP (excitation filter ET436/20 nm, emission filter ET480/40 nm), ET GFP (ET470/40 nm, ET525/50 nm), ET YFP (ET500/20 nm, ET535/30 nm), ET Chlorophyll LP (ET480/40 nm, ET610 nm LP) and ET GFP LP500 (ET470/40 nm, ET500 nm LP), used respectively for observing mTurquoise, eGFP, mVenus, chlorophyll and eGFP together with chlorophyll autofluorescence. We designed with Leica a new filter to allow for observation of RFPs like mScarlet-I without chlorophyll autofluorescence interference, the Plant RFP filter (excitation filter ET560/40 nm, emission filter ET594/10 nm).

### Laser scanning confocal microscopy

Preparation of slides: A microscope slide was fitted with a Gene Frame (ThermoFisher AB0577, 65uL, 1.5 x 1.6 cm) and a thin media pad made with 0.5x Gamborg media 1% agar. One to four Marchantia gemmae were placed on top of the pad using a small inoculation loop (ThermoFisher SL1S), a drop of sterile water placed on top of the gemmae and a #0 coverslip placed on top of the Gene Frame.

Preparation of multiwell plates: We used a 384 well plate (Greiner bio-one 781186). 0.5x Gamborg media with 1.2% (w/v) agar was poured into the selected wells for imaging, making sure they were filled up to the top and they did not contain any air bubbles inside. We filled extra sets of rows and columns of wells around our samples to reduce media evaporation. The media was allowed to solidify and leveled with a sterile scalpel if it bulged from the wells. Gene Frames (ThermoFisher AB0578, 125 uL, 1.7 x 2.8 cm) were placed to delimit the wells we were planning to image (Figure S16). One gemma was placed at the center of each filled well. The cover glass was cleaned with 70% EtOH and sprayed on one side with B-clean anti-fog spray. The anti-fog solution was let to evaporate and any residue removed with a lens tissue moving in one direction to avoid any visual artifacts. A coverslip was placed on top of the gene frame, making sure the anti-fog treated side was facing down, and the plate was ready for imaging. Gemmae could be imaged at several time points while growing inside the plate for up to 7 days.

Imaging of plants in Petri dishes: When imaging plants grown on media plates in order to follow growth processes (Figure 7, S14, S15), we let the plants grow under normal culture conditions, removed the lid for imaging and when finished, replaced the lid and returned the plants to the growth chamber.

Imaging with confocal microscopy: Images were acquired on a Leica SP8X Spectral Confocal Microscope upright system equipped with a 460-670 nm super continuum white light laser, 2 CW laser lines 405nm, and 442 nm, and 5 Channel Spectral Scanhead (4 hybrid detectors and 1 PMT). For slides, imaging was conducted using either a 10x air objective (HC PL APO 10x/0.40 CS2), a 20x air objective (HC PL APO 20x/0.75 CS2) or a 40x water immersion objective (HC PL APO 40x/1.10 W CORR CS2). For plants on multiwell plates or on plates without lid, 10x or 20x objectives could be used. Imaging settings varied depending on number of channels. When observing more than one fluorescent protein, sequential scanning mode was selected. Excitation laser wavelength and captured emitted fluorescence wavelength window were as follows: for mTurquoise2 (442 nm, 460-485 nm), for eGFP (488 nm, 498-516 nm), for mVenus (515 nm, 522-540 nm), for mScarletI (569 nm, 585-610 nm) and for chlorophyll autofluorescence (488 or 515, 670-700 nm). Chlorophyll autofluorescence was imaged simultaneously with eGFP or mVenus.

### Registries and Metadata

Benchling was used as DNA editor and registry of constructs. The Benchling Golden Gate assembly wizard allows for building of type IIS DNA constructs keeping the lineage information, list and order of plasmids used to assemble the final one, like all the L0 plasmids used to make a L1 plasmid (https://benchling.com/tutorials/11/golden-gate-assembly). Curation of parts is facilitated by the auto-annotate sequence tool(https://benchling.com/tutorials/34/auto-annotate). The bioregistry (https://benchling.com/tutorials/47/setting-up-your-bioregistry) option allows establishing of registries for different type of parts with specific associated metadata (schema), which facilitates standardization (https://help.benchling.com/articles/2725066-configure-your-registry). All parts of a registry, with its unique identifier for each part and all associated metadata can be exported in bulk as a csv file (https://help.benchling.com/articles/2725575-export-registry-data-in-bulk), and we use R studio and R packages tidyr and ggplot2 to edit and further process them (https://www.rstudio.com/). The plasmids in the OpenPlant are shared through a public benchling folder (https://benchling.com/susana/f_/1ptxoMWP-openplant-kit/).

## Supporting information

Supplemental Registry of parts

Supplemental RNA seq data

## ACKNOWLEDGMENTS

We thank Suvi Honkanen for sharing constructs to test as constitutive promoters. We thank Connor Tansley for sharing his experience on cryopreservation protocol. We thank Emanuele Orsini and Michelle Lim for their contribution as summer students. We thank previous and present members of the lab working with Marchantia, Nuri Purswani, Christian Boehm, Owen Male, Liat Adler, Alicja Szałapak, Lukas Mueller, Alan Marron, Jenna Rever and Bernardo Pollak; the latter especially for his doctoral thesis ^63^ that described a plate-based transformation method, and a Nop1-targeted gRNA that provided a working design for this work. We thank the Cambridge Marchantia community, specially Chiara Airoldi, Giulia Arsuffi and Philip Carella for useful discussions and sharing of materials, and the ROC (Researchers with OpenPlant in Cambridge) for useful discussions on standardization and sharing tools. We thank Benchling for diligent support in implementing Registries. We thank Plant Sciences Department facilities and administrative staff for their support. This work was funded as part of the BBSRC/EPSRC OpenPlant Synthetic Biology Research Centre Grant BB/L014130/1 to NJP and JH, BBSRC BB/F011458/1 for confocal microscopy to JH, BBSRC Research Studentships for MR (1943399) and MD (1497818), and BBS/OS/GC/ 000013C Synthetic Biology Tools and Resources for genetic Improvement in the Developing World and BBS/E/T/000PR9815A Earlham BioFoundry, a BBSRC-funded National Capability for NJP.

## SUPPORTING INFORMATION

Detailed experimental protocols can be found at https://www.protocols.io/groups/openplant-project

Supplementary-FigureS1-S16-TableS1-S2. Supplementary Figures and Tables to describe experiments and Materials and Methods.

Registry_OpenPlantkit.csv File with registry of OpenPlant kit plasmids, with sequences and metadata.

MpRNAseq.xlsx Spreadsheet file with all publicly available RNAseq experiments reanalyzed, expression levels for all Marchantia genes and TPM stats.

---

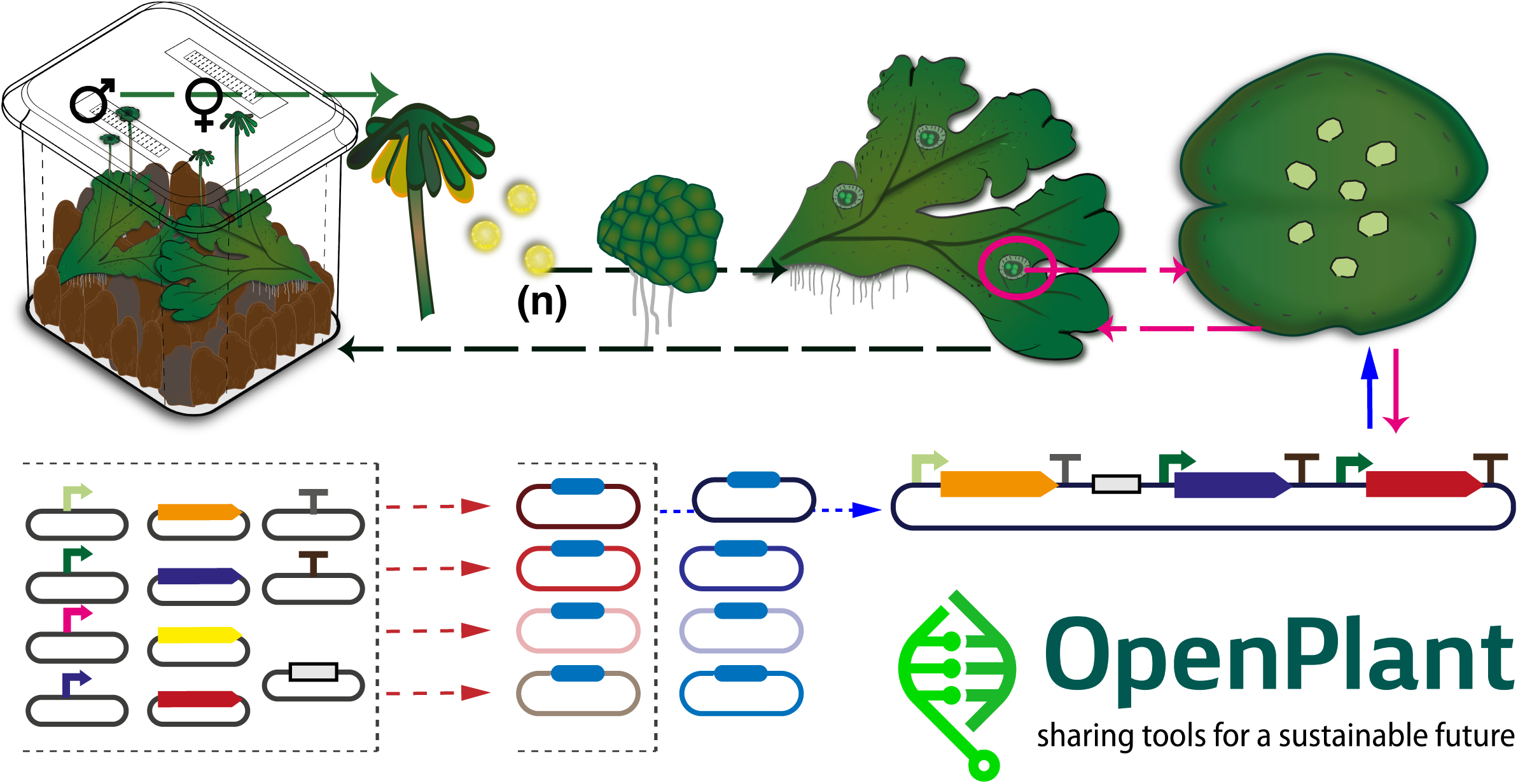

