## Supplemental Figures for "Systematic tools for reprogramming plant gene expression in a simple model, *Marchantia polymorpha*"

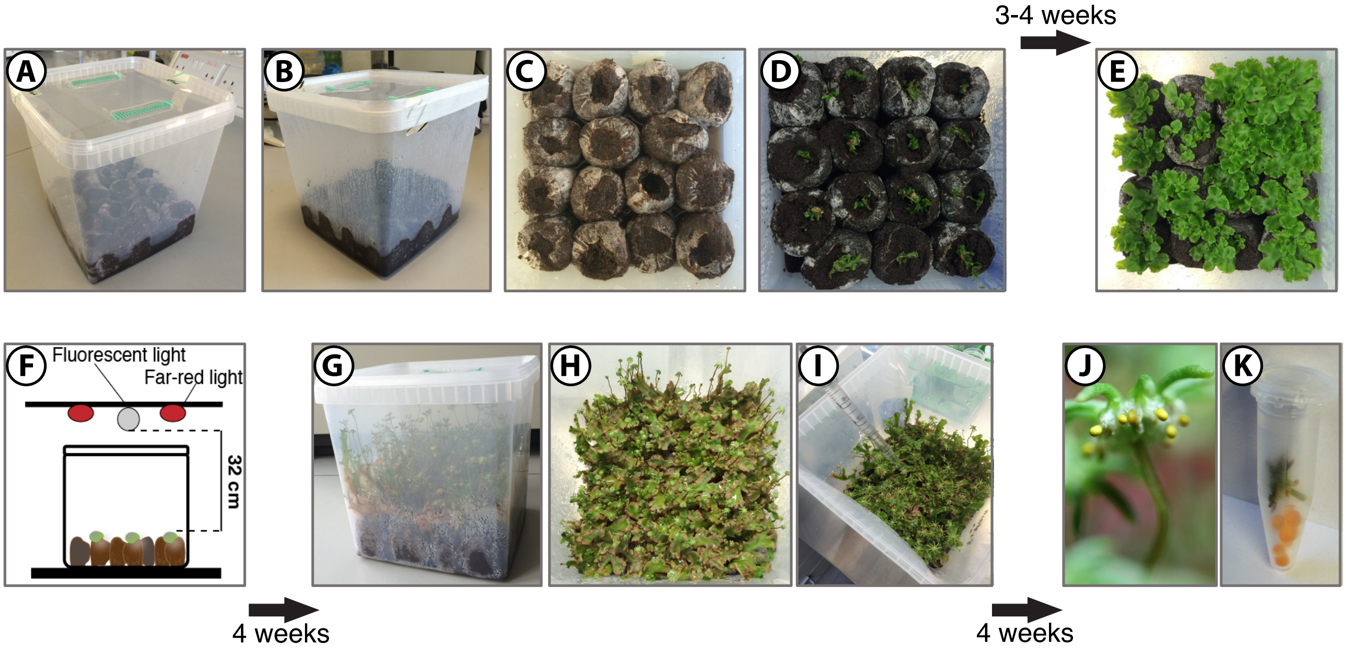


**Figure S1.** Techniques for axenic propagation of Marchantia spores. (A-B) 15 Jiffy-7 compressed peat disks were placed in a TP5000+TPD5000 Microbox with #40 green filter, 800 mL of water added, the lid was closed and the Microbox was autoclaved. (C-D) After autoclaving all work was performed in a laminar flow hood. One 10 mm x 10 mm thallus fragment or 3 gemmae were placed on each hydrated Jiffy-7 pellet using sterile tweezers. Another 150-200 mL of sterile water were added into the Microbox, the lid was sealed and the Microbox was placed at 21 °C under continuous light 150 μE. (E-F) After one month, Microboxes were transferred to long day conditions (16 hours light / 8 hours dark) with light intensity of 150 μE supplemented with far-red light (Philips GreenPower LED HF far red, peak emission around 730-740 nm, at a distance from plants of 32 cm). (G-H) In about 4 weeks, the plants produced mature male and female reproductive organs. (I) In a flow hood, the lid of the Microbox was opened, and using a pipette, 50-100 μL of sterile water was added to the top of male reproductive organs. After a few seconds the sperm was visible as a cloudy exudate in the water drop. For fertilization, the drop was transferred, with a disposable sterile plastic pipette, to the top of a female reproductive organ. (J) After one more month, mature yellow colored sporangia were visible and ready for collection. In one Microbox, 60-70 fertile female gametophores usually developed, producing several hundred mature sporangia. (K) 1-2 mature sporangia were placed in a 1.5 mL centrifuge tube with 5 silica gel beads and stored at 4 °C if the spores were to be used within 1-2 months. For longer storage, sporangia were allowed to desiccate for a week at 4 °C and then stored at -80 °C.

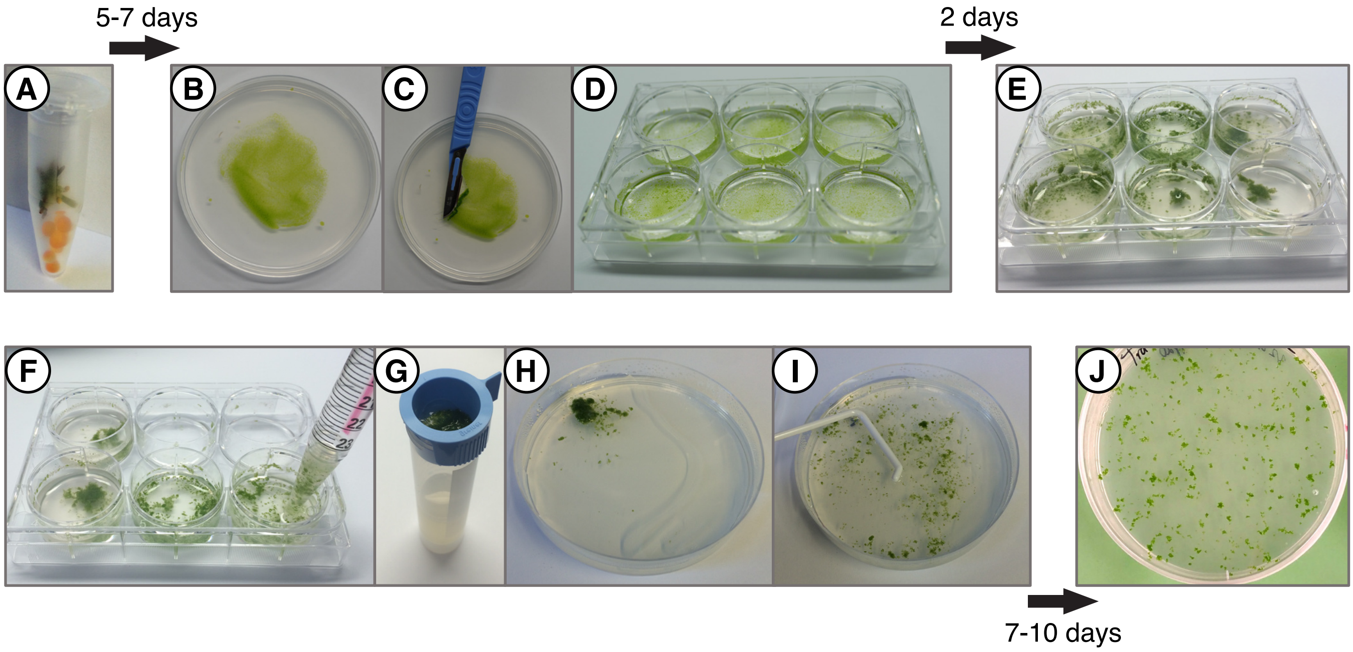


**Figure S2.** Agrobacterium-mediated transformation in multi-well plates. (A) Marchantia spores were sterilised and plated on 0.5x Gamborg B5 plates. (B-D) After 5-7 days, sporelings (germinated spores) were transferred into a 6-well plate containing 4 ml per well of 0.5x Gamborg B5 media plus supplements and 100 µM acetosyringone, using a sterile scalpel. 100 µL of an Agrobacterium culture transformed with a plasmid of interest was added to each well. (E) The plate was then placed on a shaker for 2 days at 21°C and continuous light. (F-G) After co-cultivation of sporelings and Agrobacterium, sporelings in each well were washed with 50 mL of sterile water on a cell strainer and (H-I) then plated on 0.5x Gamborg B5 plates with suitable antibiotics. (J) After 7-10 days, successful transformants were visible on the plate and could be screened with fluorescent microscopy.

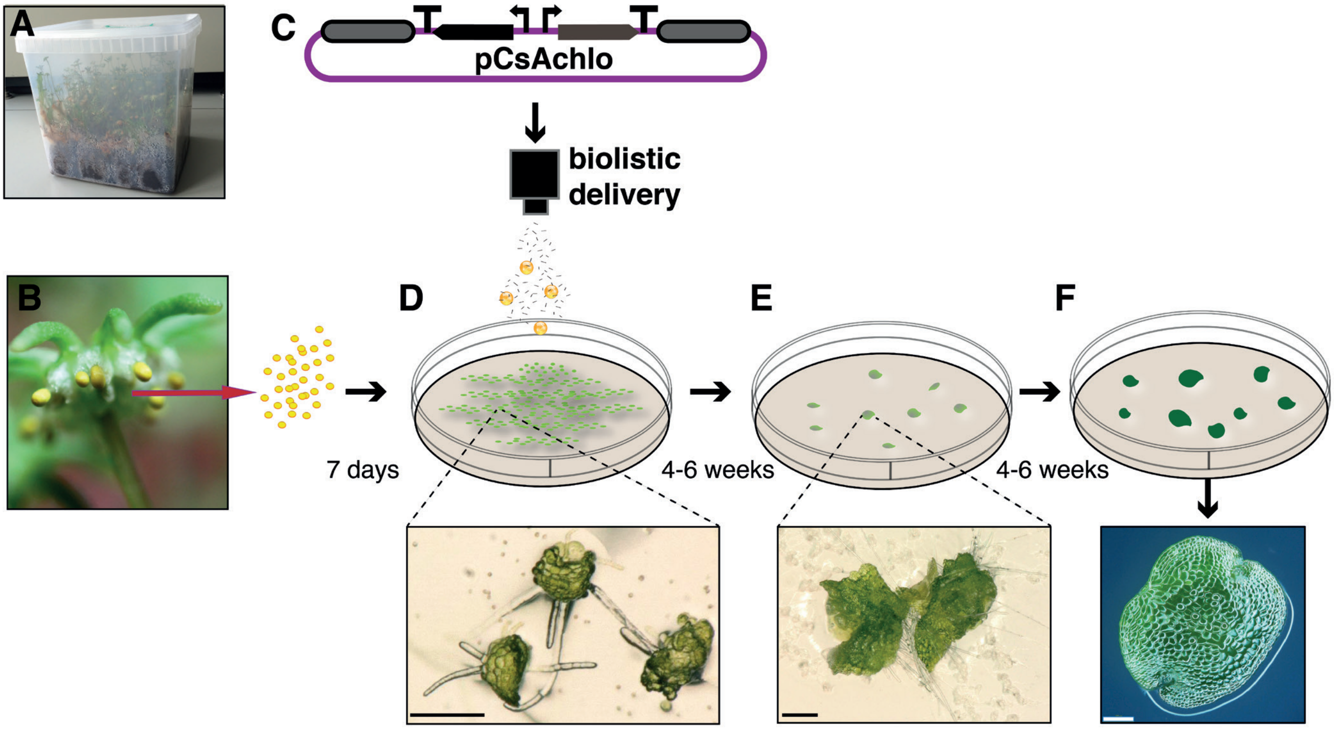


**Figure S3.** Outline of the chloroplast transformation method. (A) Microboxes were used to produce spores (Figure S1). (B) Spores were plated on 0.5x Gamborg B5 plates and incubated under continuous light for 7 days. (C-D) 7 day old sporelings were bombarded with DNAdelTM nanoparticles coated with the plasmid DNA of interest and placed in a growth room for 2 days. (E) They were then transferred to 0.5x Gamborg B5 plates with spectinomycin (500 µg/mL), and successful transformants were visible after 4-6 weeks. (F) Transformants were then transferred onto a fresh 0.5x Gamborg B5 plate containing spectinomycin. After 4-6 weeks in this second round of selection, plants produced gemmae cups, and gemmae could be tested for genotype and homoplasmy by PCR. Scale bar: (D-E) 200 µm, (F) 100 µm.

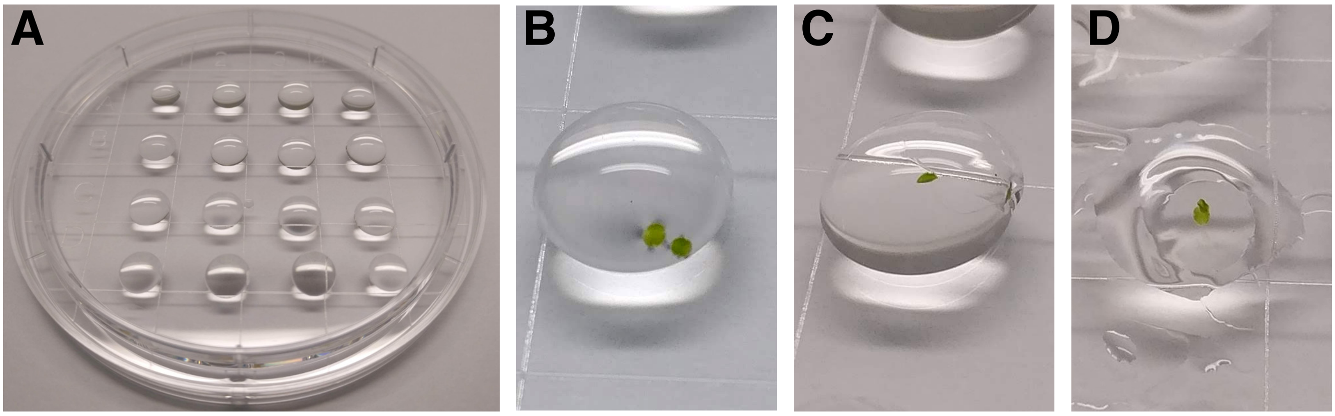


**Figure S4.** Cryopreservation of Marchantia gemmae. (A) Drops of alginate solution were prepared on a Petri dish, and (B) up to five Marchantia gemmae were transferred inside the alginate beads. (C) Beads polymerized upon addition of calcium chloride solution to the top of the bead, and gemmae were immobilized. (D) Gemmae were then soaked in dehydration solution and air-dried for at least 2 h. The dried beads with gemmae inside were placed in 1.5 mL Eppendorf tubes, frozen and kept at -80 °C.

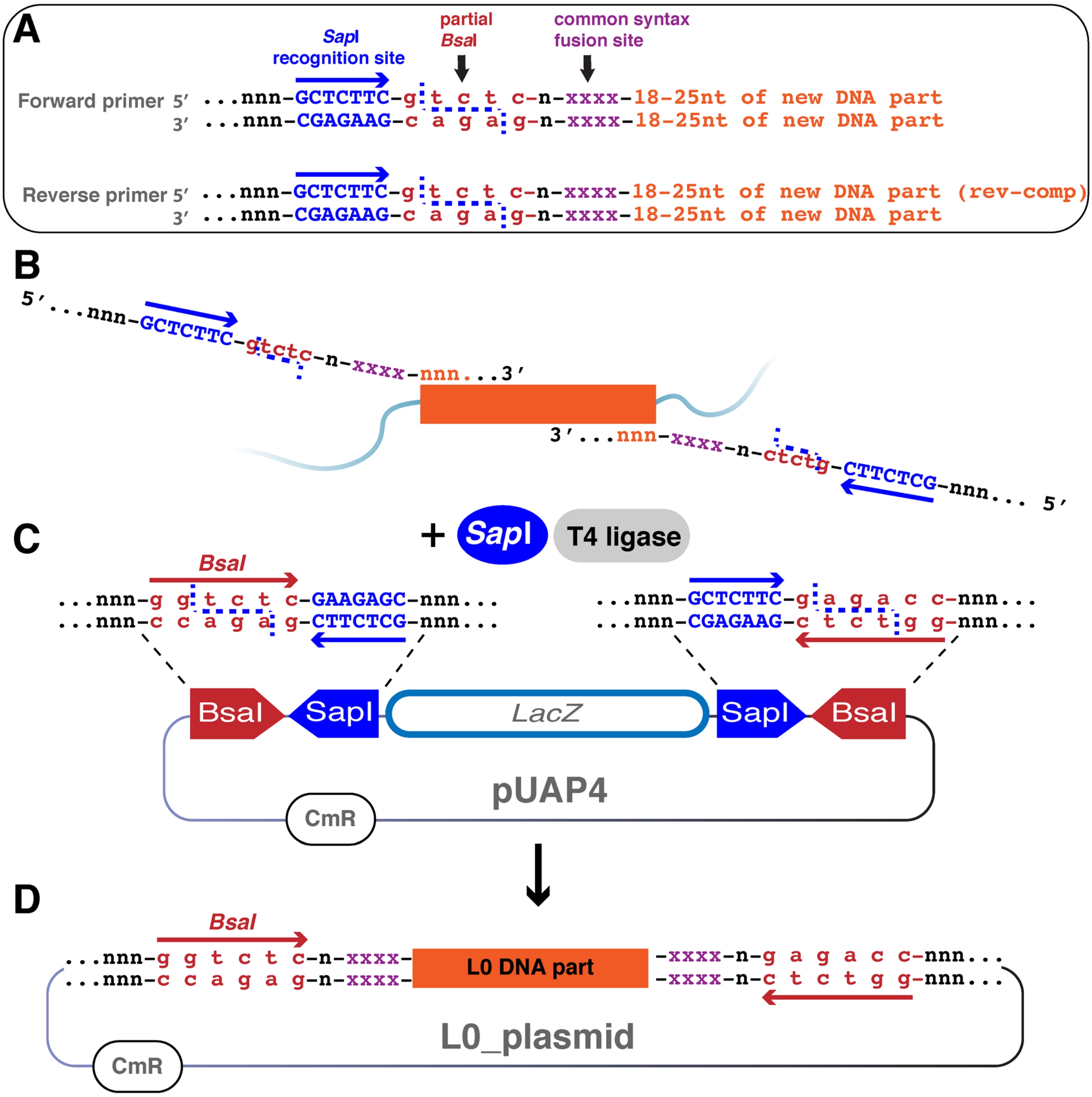


**Figure S5.** Design of primers for cloning of Level 0 parts into pUAP4 vectors. (A) To clone a standardized L0 part, 5' and 3' common syntax sites were fused to the DNA part by PCR, using specially designed primers. Primers included the sequences: 3 random bp (in black), the 7 bp *Sap*I recognition site (in blue), 5 bp that correspond to a partial *Bsa*I recognition site (in red), one random bp for spacing (in black), and the common syntax fusion site (in purple) followed by 18-25 bp complementary to the DNA part to be amplified (in orange). (B) DNA parts were amplified with a high fidelity DNA polymerase and (C) cloned into pUAP4 using *Sap*I type IIS assembly to produce (D) the desired Level 0 plasmid. Once cloned into pUAP4, the full *Bsa*I recognition site is reconstituted. *Bsa*I type IIS cloning can be used to assemble multiple L0 parts into a transcription unit (L1). Blue arrows: *Sap*I recognition site. Blue dashed lines: *Sap*I cleavage site. Red arrows: *Bsa*I recognition site. CmR: chloramphenicol bacterial resistance cassette. LacZ: lacZα cassette for blue-white screening of colonies (negative blue colonies contain undigested pUAP4, positive white colonies contain a L0 part inserted into pUAP4).

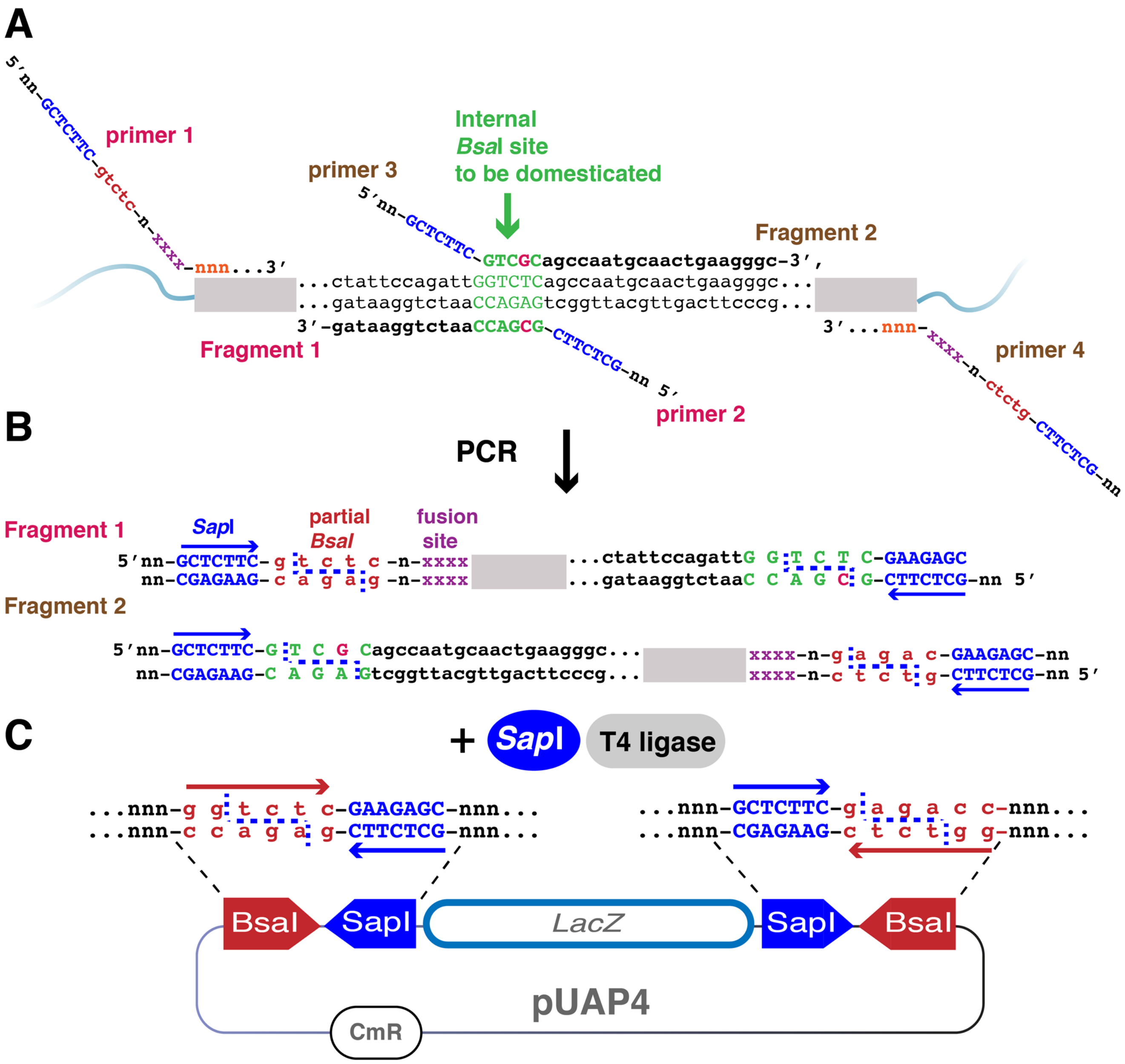


**Figure S6.** Guide to domestication of new DNA parts. Removal of internal *Bsa*I and *Sap*I sites is necessary when a DNA sequence of interest contains an internal *Bsa*I or *Sap*I recognition site. Overlapping PCR reactions can be used to mutate unwanted sites. Two PCR fragments are amplified upstream (fragment 1, with primers 1 and 2) and downstream (fragment 2, with primers 3 and 4) of the unwanted recognition site. Both primer 2 and 3 can be designed with a single nucleotide mismatch to alter the target sequence (taking care to not alter amino acid composition if the region is in the coding sequence). Primers also contain *Sap*I recognition sites to allow the two fragments to be ligated together into pUAP4 using *Sap*I type IIS assembly. Primer 1 and primer 4 are designed according to the guidelines for cloning of L0 parts into pUAP4 (see Figure S5). Blue arrows: *Sap*I recognition site. Blue dashed lines: *Sap*I cleavage site. Red arrows: *Bsa*I recognition site. CmR: chloramphenicol bacterial resistance cassette. LacZ: lacZα cassette for blue-white screening of colonies.

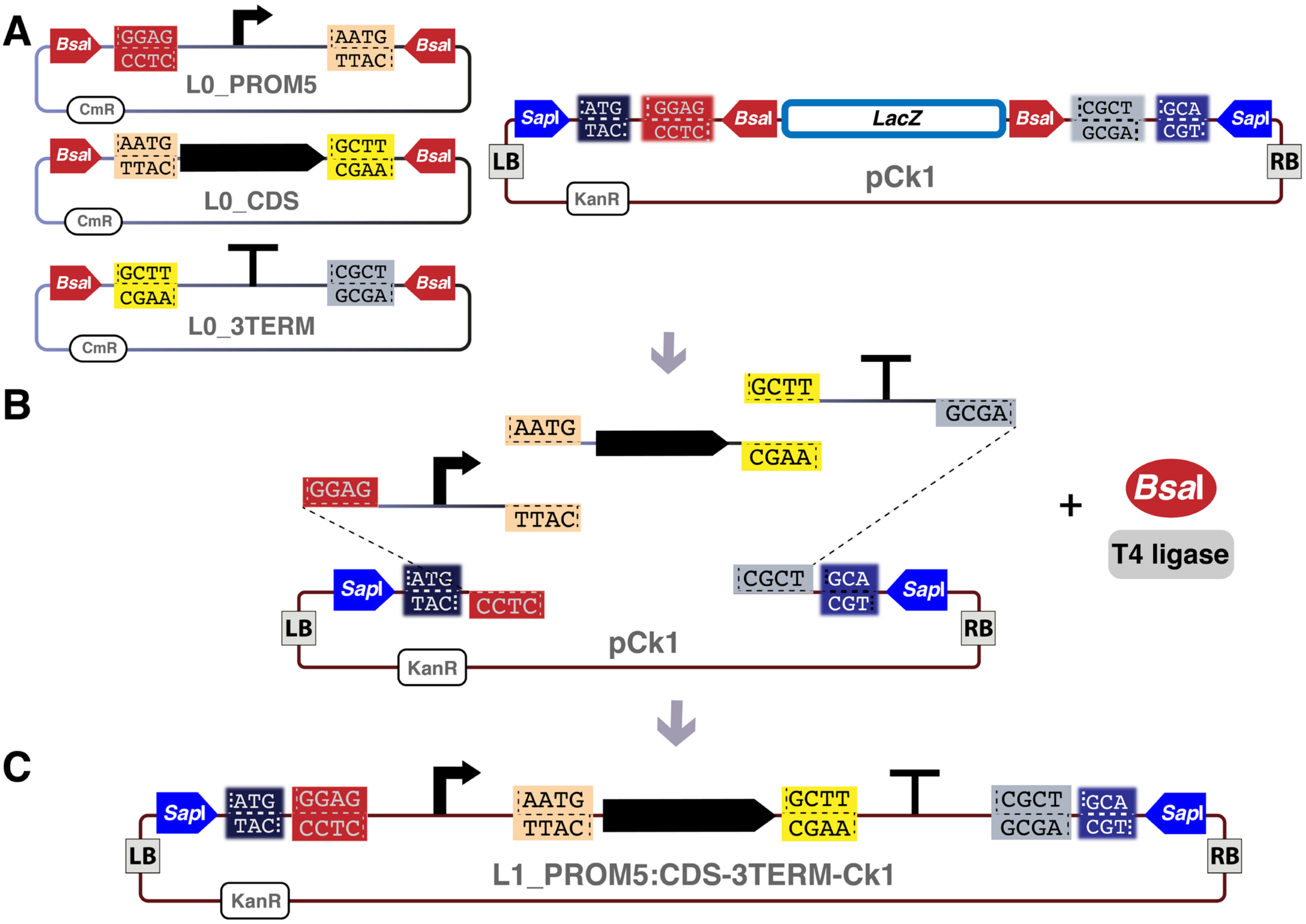


**Figure S7.** Assembly of multiple Level 0 parts into a transcription unit. (A-C) In a single-step digestion-ligation reaction, plasmids containing standard L0 parts (e.g. PROM5, CDS and 3TERM) are mixed with a pCk plasmid (e.g. pCk1), *Bsa*I and T4 ligase enzymes. (A) L0 plasmids contain convergent *Bsa*I sites and pCk plasmids contain divergent *Bsa*I sites. (B) Digestion of a L0 plasmid with *Bsa*I releases the part with 4 bp overhangs (e.g. GGAG) as sites for fusion according to the common syntax. Similarly, digestion of a pCk plasmid with *Bsa*I releases the LacZ cassette and reveals the acceptor fusion sites as overhangs in the backbone (example CTCC). (C) L0 parts are ligated into the pCk vector with the fusion sites enabling directional assembly of DNA parts to build the desired L1 (in this case the L1_PROM5:CDS-3TERM-Ck1). Once the L1 has been assembled, it does not contain *Bsa*I sites and thus cannot be further digested. After transformation into competent *E. coli* cells and plating on selection media with kanamycin, the colonies with undigested L0 plasmids will not be selected due to their different antibiotic resistance cassette (chloramphenicol) and colonies with undigested pCk plasmids will be blue in plates containing X-gal. Red arrows: *Bsa*I restriction sites. Blue arrows: *Sap*I restriction sites. CmR: chloramphenicol bacterial resistance cassette. KanR: kanamycin bacterial resistance cassette. LacZ: lacZα cassette for blue-white screening of colonies.

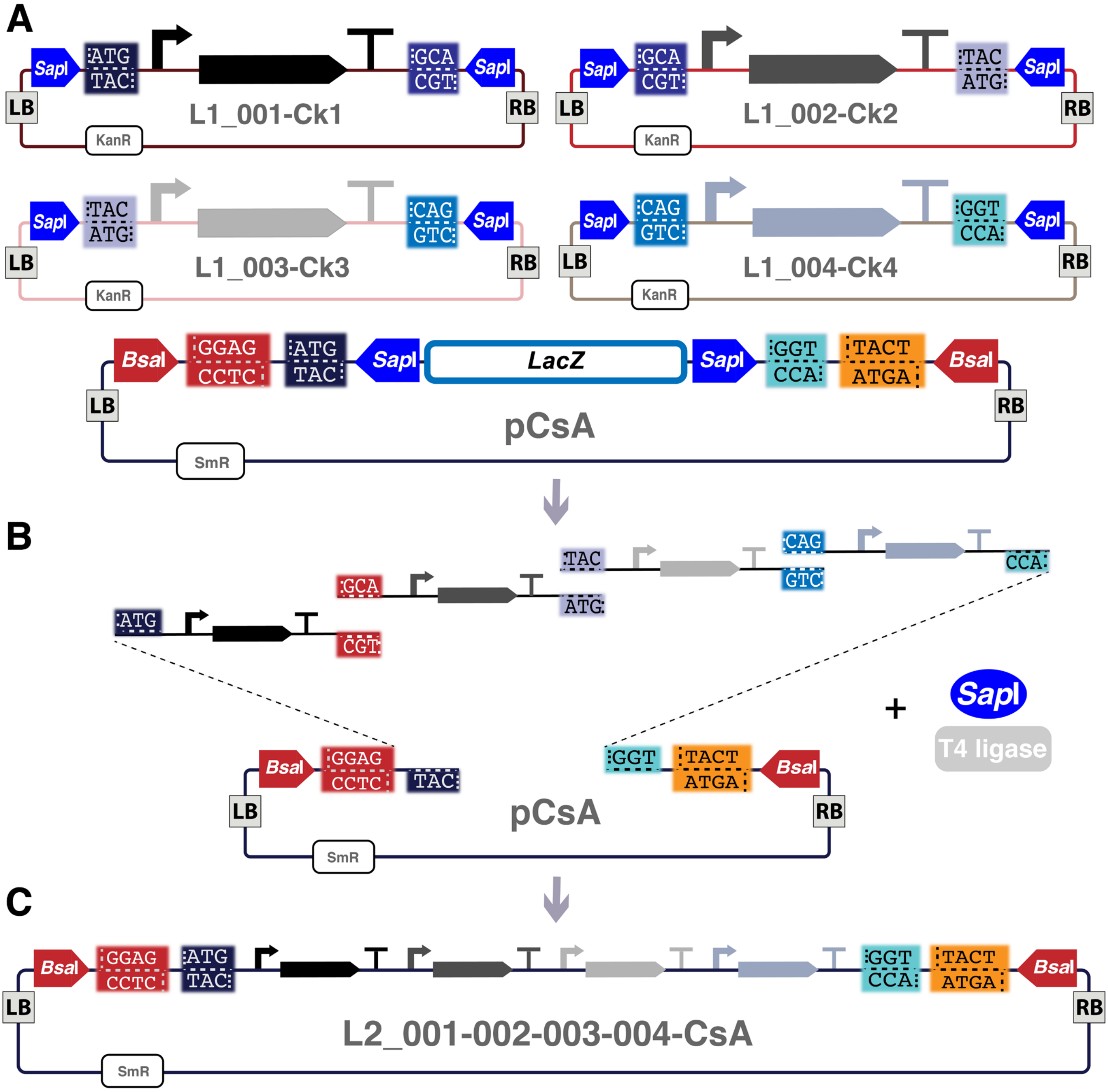


**Figure S8.** Loop assembly of four transcription units into a Level 2 plasmid. (A-C) In a one-step reaction, four transcription units assembled in pCk vectors (in this case L1_001-Ck1, L1_002-Ck2, L1_003-Ck3, L1_004-Ck4) were mixed with a pCs plasmid (in this case, pCsA), *Sap*I and T4 ligase enzymes. (A) L1 plasmids contain convergent *Sap*I sites and pCs plasmids contain divergent *Sap*I sites. (B) Digestion of a L1 plasmid with *Sap*I releases the L1 construct with its specific 3 bp Loop fusion sites as overhangs (e.g. ATG). Similarly, digestion of a pCs plasmid with *Sap*I releases the LacZ cassette and reveals the acceptor 3 bp fusion sites as overhangs in the backbone (example CAT). (C) Released L1 fragments are ligated into the pCs vector, with the fusion sites enabling directional assembly of DNA parts and building of the desired L2 device (in this case the L2_001-002-003-004-CsA). Once the L2 has been assembled, it lacks *Sap*I sites and thus cannot be further digested. The L2 reaction was transformed into competent *E. coli* cells and plated on selection media with spectinomycin. Undigested L1 plasmids do not carry the correct antibiotic resistance cassette and undigested pCs plasmids will be marked blue in plates containing X-gal. Red arrows: *Bsa*I restriction sites. Blue arrows: *Sap*I restriction sites. KanR: kanamycin bacterial resistance cassette. SmR: spectinomycin bacterial resistance cassette. LacZ: lacZα cassette for blue-white screening of colonies.

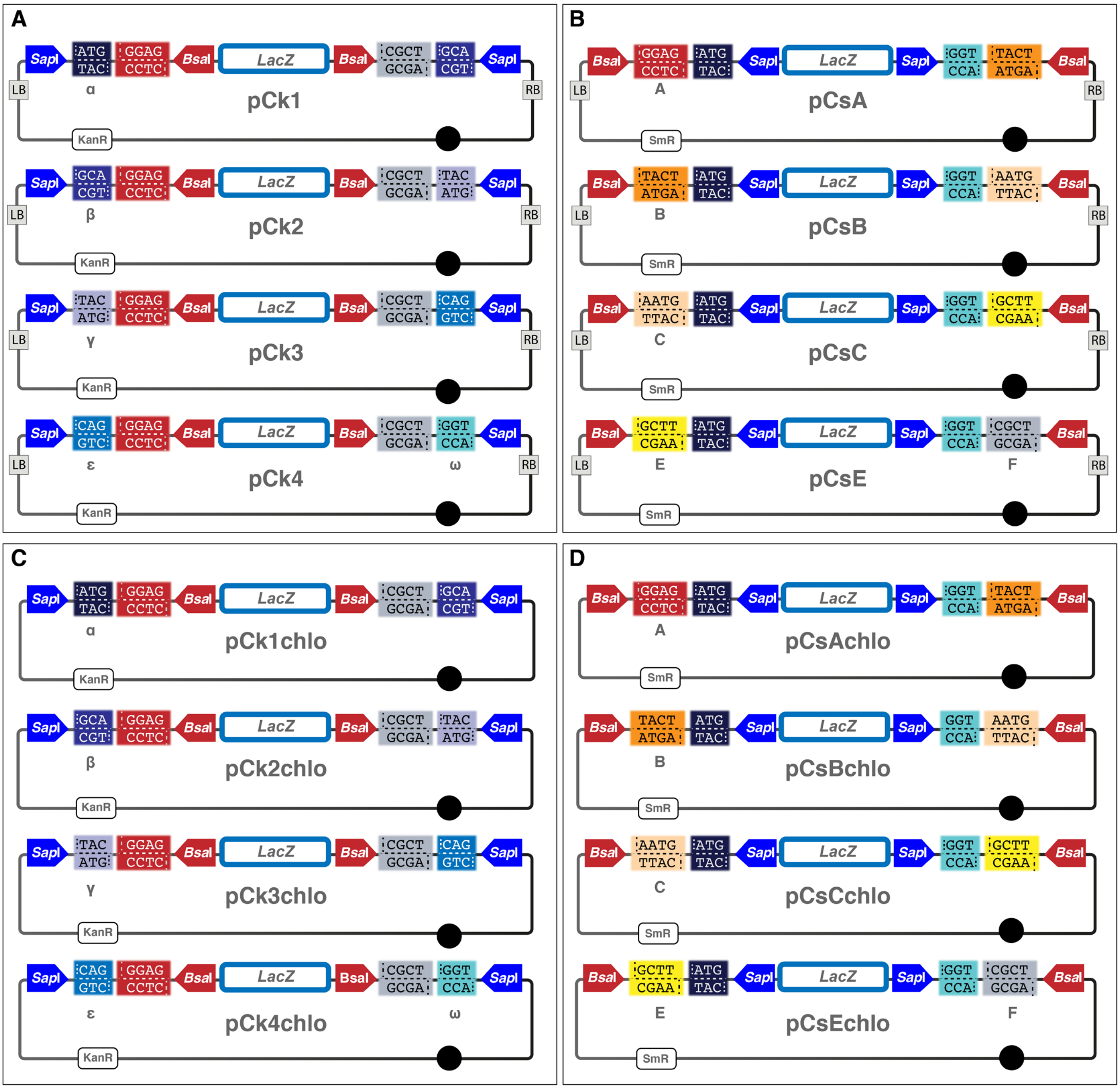


**Figure S9.** Improved Loop vectors for nuclear and chloroplast transformation. (A) pCk vectors (pCk1, pCk2, pCk3 and pCk4) all contain divergent *Bsa*I restriction sites (red arrows) adjacent to specific Loop fusion sites [A (GGAG) and F (CGCT)] to allow assembly of different L0 parts into a L1 construct using a pCk vector and *Bsa*I. They also contain convergent *Sap*I restriction sites (blue arrows) next to specific Loop fusion sites to allow assembly different L1 constructs into a L2 construct using a pCs vector and *Sap*I. The fusion sites are: α (ATG), β (GCA), δ (TAC), ε (CAG) and ω (GCT), as pCk1 α-β, pCk2 β-δ, pCk3 δ-ε and pCk4 ε-ω. (B) pCs vectors (pCsA, pCsB, pCsC and pCcE) all contain divergent *Sap*I restriction sites (blue arrows) adjacent to specific Loop fusion sites, α-ω, to assemble up to four L1 constructs into a L2 device using a pCs vector and *Sap*I. They also contain convergent *Bsa*I restriction sites (red arrows) next to specific Loop fusion sites to allow assembly of up to four L2 devices into a L3 device using a pCk vector and *Bsa*I. The fusion sites are: A (GGAG), B (TACT), C (AATG), E (GCTT) and F (CGCT), which are present as: pCs1 A-B, pCs2 B-C, pCs3 C-E, pCs4 E-F. (C-D) Transformation vectors for chloroplast applications, pCkchlo vectors (C) and pCschlo vectors (D), contain, respectively, same *Bsa*I, *Sap*I restriction sites, and fusion sites as the pCk and pCs sets, but they do not contain the left (LB) and right border (RB) repeats from the nopaline C58 T-DNA for Agrobacterium-mediated nuclear transformation. KanR: kanamycin bacterial resistance cassette. SmR: spectinomycin bacterial resistance cassette. LacZ: lacZα cassette for blue-white screening of colonies.

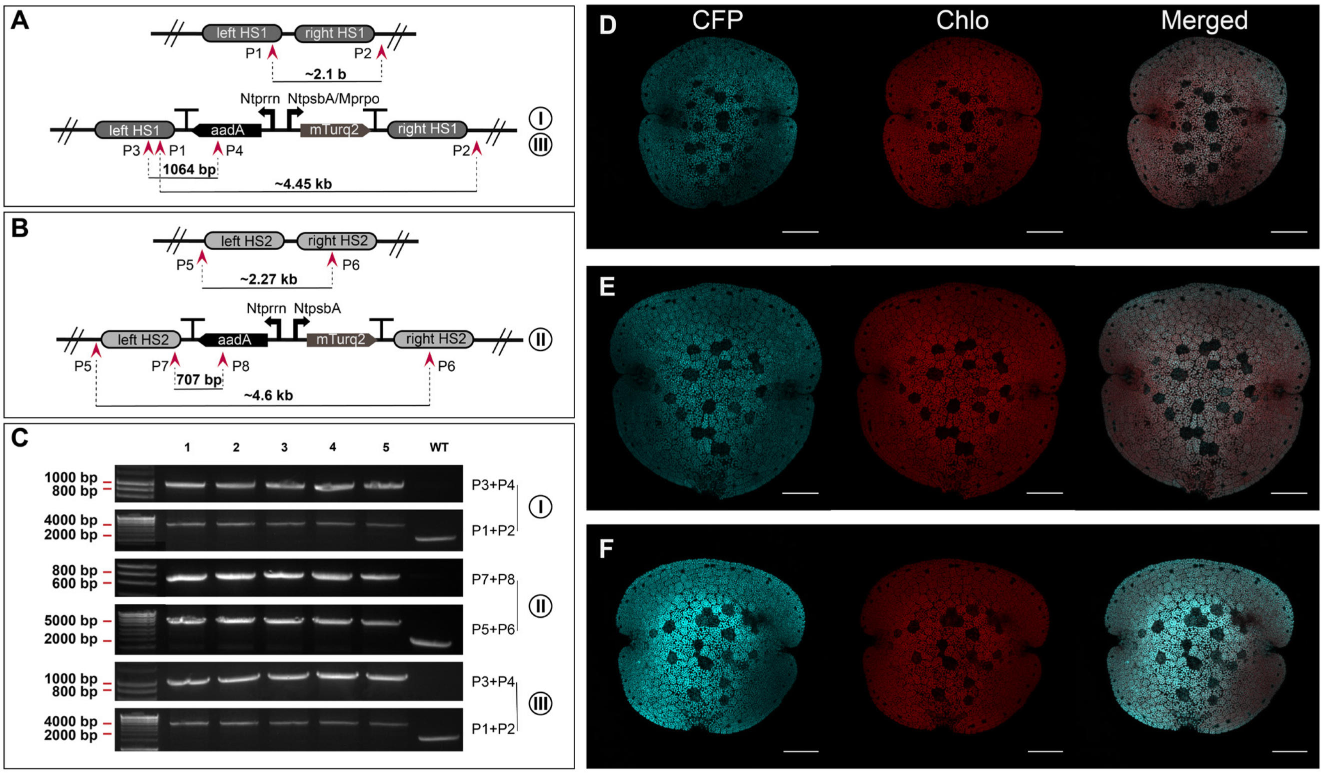


**Figure S10**. Validation of DNA parts and vectors for chloroplast transformation. (A) Schematic representation of the *trnG–trnfM* target region (flanked by HS1-L and HS1-R) in the wild type chloroplast genome (top) and the same region after integration of the DNA construct I or III (bottom). Construct **I**: p5-NtpsbA:mTurquoise2_chlo for integration into intergenic region 1. Construct **III**: p5-Mprpo:mTurquoise2_chlo for integration into intergenic region 1. (B) Schematic representation of the *rbcL-trnR* target region (flanked by HS2-L and HS2-R) in the wild type chloroplast genome (top) and the same region after integration of the DNA construct II (bottom). Construct **II**: p5-NtpsbA:mTurquoise2_chlo for integration into intergenic region 2. (A-B) Red arrows indicate the position of the PCR primers used for the detection of wild type or homoplasmic transplastomic lines. Maps not in scale. (C) PCR analysis of genomic DNA isolated from wild type and transplastomic plants. Integrity of the reporter gene (top, primers P3+P4 or P7+P8) and homoplasmy (bottom, primers P1+P2 or P5+P6) were confirmed for transplastomic lines after 2 months of subculture under selective conditions. The primer pair used for each PCR are shown next to the gel images. (D-F): Images of 0 days transplastomic gemmae expressing (D) construct I, (E) II or (F) III. (D-F) Left: mTurquoise2cp, middle: Chlorophyll autofluorescence, right: merged images. Scale bar: 100 µm.

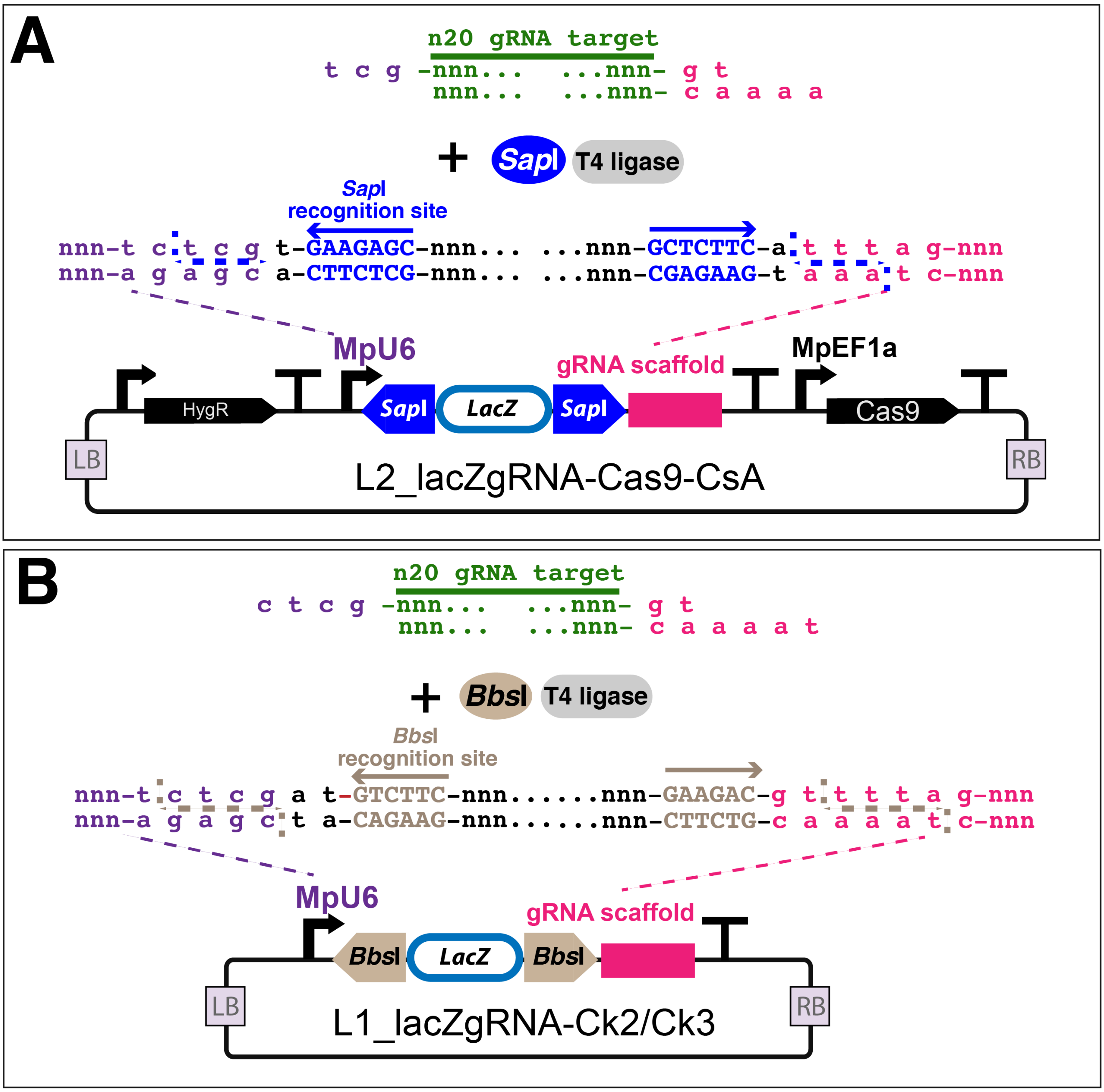


**Figure S11**. Design of oligonucleotide primers for rapid cloning of gRNAs. (A) Design of synthetic oligonucleotides for gRNA *Sap*I mediated cloning into the L2_lacZgRNA-Cas9-CsA vector. The L2_lacZgRNA-Cas9-CsA *Sap*I digested vector has AGC and TTT overhangs. Therefore, oligos for gRNA should be designed such that the forward strand has a 5' overhang of TCG and the reverse strand has a 5' overhang of gt-AAA (addition of “gt” nucleotides is necessary to reconstitute the full sequence of the gRNA scaffold in pink). Blue arrows: *Sap*I recognition site. Blue dashed lines: *Sap*I cleavage site. (B) Design of oligos for gRNA *Bbs*I mediated cloning into L1_lacZgRNA-Ck2 or L1_lacZgRNA-Ck3 vectors. The L1_lacZgRNA-Ck2/3 *Bbs*I digested vectors have GAGC and TTTA overhangs. Synthetic oligos for the gRNA should be designed so that the forward strand has a 5' overhang of CTCG and the reverse strand has a 5' overhang of gt-AAAT (addition of “gt” nucleotides is necessary to reconstitute the full sequence of the gRNA scaffold in pink). Light brown arrows: *Bbs*I recognition site. Light brown dashed lines: *Bbs*I cleavage site. (A-B) LacZ: lacZα cassette for blue-white screening of colonies.

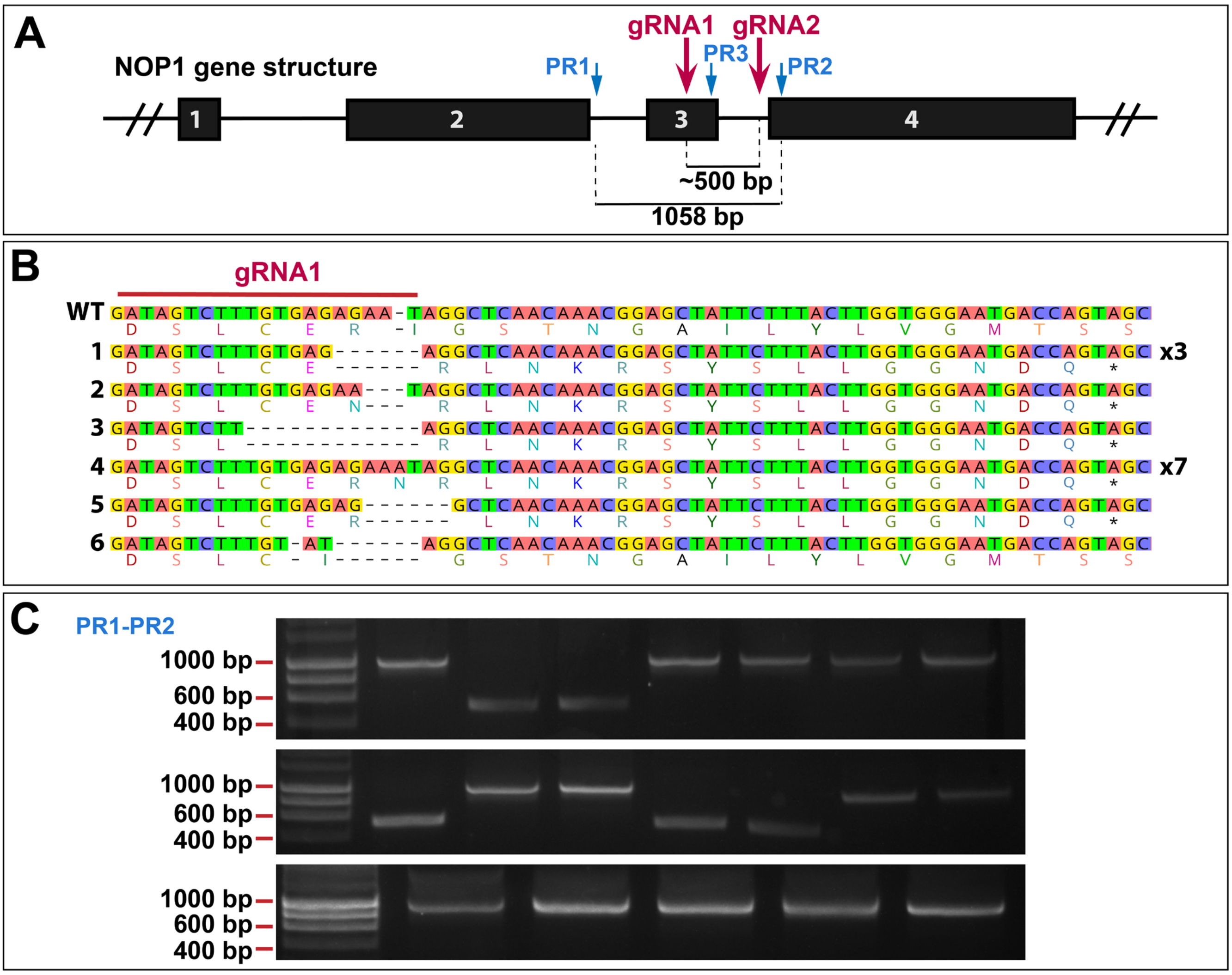


**Figure S12**. Cas9-CRISPR mediated gene editing and generation of deletions. (A) Schematic representation of the Mp*NOP1* gene structure showing exons as black boxes. The gRNAs for CRISPR and primers for PCR-based genotyping are shown in red and blue, respectively. (B) Sequence analysis of *nop1* knockout mutant lines obtained with the L2 gRNA1 CRISPR vector (L2_NOP1gRNA-Cas9-CsA, Figure 4A). The wild type Marchantia Cam-1 sequence is shown at the top, with the 20 bp gRNA1 target sequence highlighted with a red line. The amino acid sequence is depicted below the nucleotide sequence. Primers used: PR1-PR3. Of the 14 mutants analyzed, sequence deletions of identical to line 1 occured 3 times and for line 4, 7 times. (C) Gel electrophoresis images of PCR-based genotyping of the mutants obtained with the L2 gRNA1-gRNA2 CRISPR vector (L2_2xNOP1gRNA-Cas9-CsA, Figure 4C). Primers used: PR1-PR2. Expected size of PCR products for wild type or mutants with small deletions (as in B) is approximately 1000 bp. Expected size of PCR products when large deletions occurred is approximately 500 bp. The large deletions were further confirmed by Sanger sequencing.

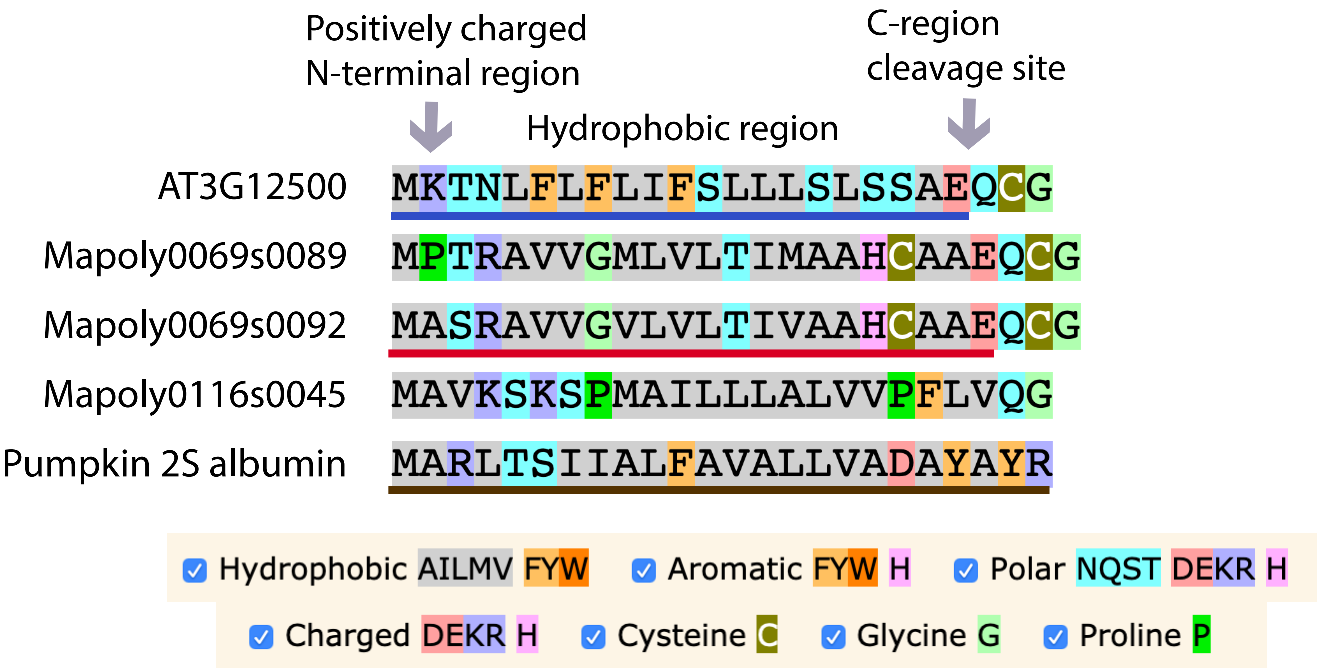


**Figure S13.** Alignment of ER signal peptides. ER signal peptides do not show conserved primary amino acid sequence, but typically include a positively charged N-terminal region, followed by a hydrophobic region, and C-region that defines the cleavage site [1]. The ER signal peptide from the Arabidopsis basic chitinase (AT3G12500) (underlined in blue) has been used in other plant species to confer ER localization. The ER signal peptide from pumpkin 2S albumin (underlined in brown) has been shown to work in Marchantia. In this work, we identified three predicted native chitinases in Marchantia (Mapoly0069s0089, Mapoly0069s0092, and Mapoly0116s0045), tested the leader peptide from one of them, Mapoly0069s0092 (underlined in red), and confirmed that it conferred ER localization in Marchantia. Protein alignment was performed using MSAReveal.org

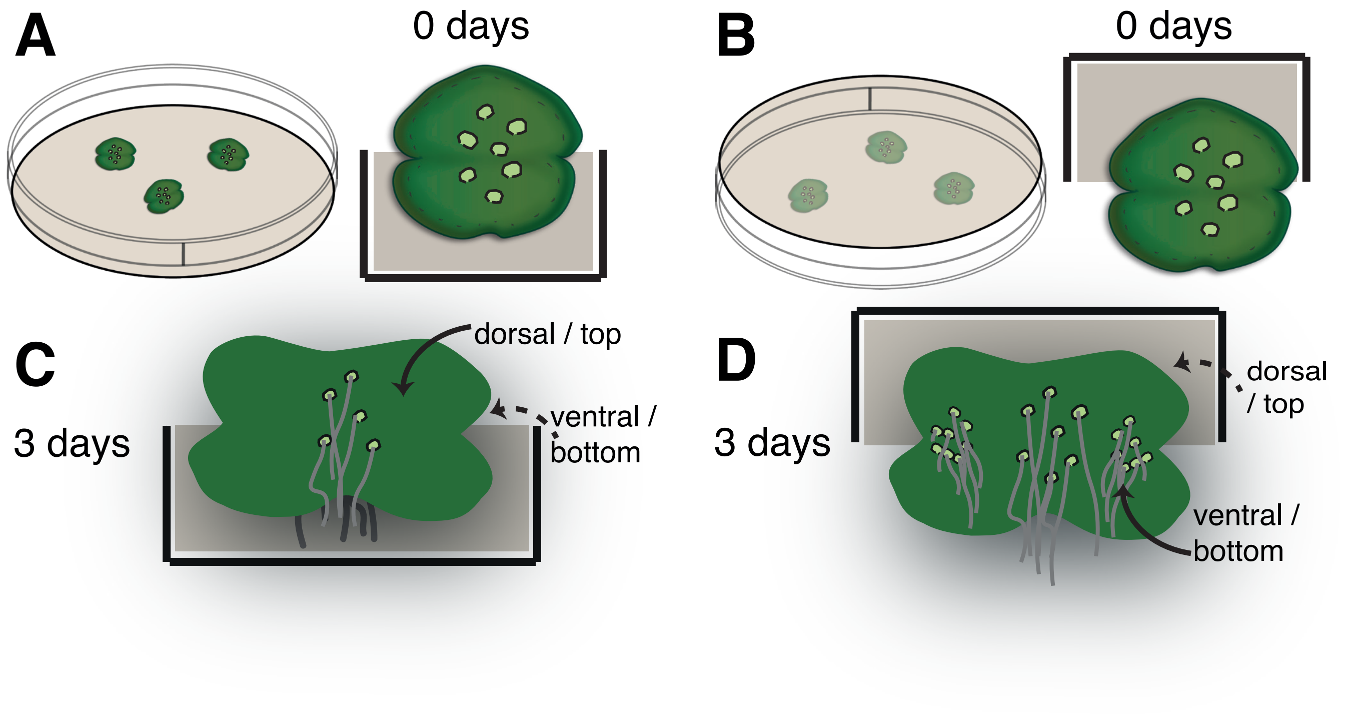


**Figure S14**. Schematic representation of gemmae growing in normal and inverted plates. The bottom side of gemmae acquires ventral identity both in normal and inverted plates. Gemmae removed the gemma cups (0 day) have rhizoid precursors present on both sides of the central part of the gemmae. Rhizoids are larger size cells that lack mature chloroplasts (light green cells in A,B). They elongate once gemmae are placed on media and dormancy is released (C,D). As gemmae grow, they acquire dorso-ventral polarity, with the dorsal (top) side developing air chambers and the ventral (bottom) side developing predominant rhizoids (A,C). If plates are inverted (B), the lower sides of the inverted gemmae form new rhizoids and adopt ventral identities, despite not touching the media (D).

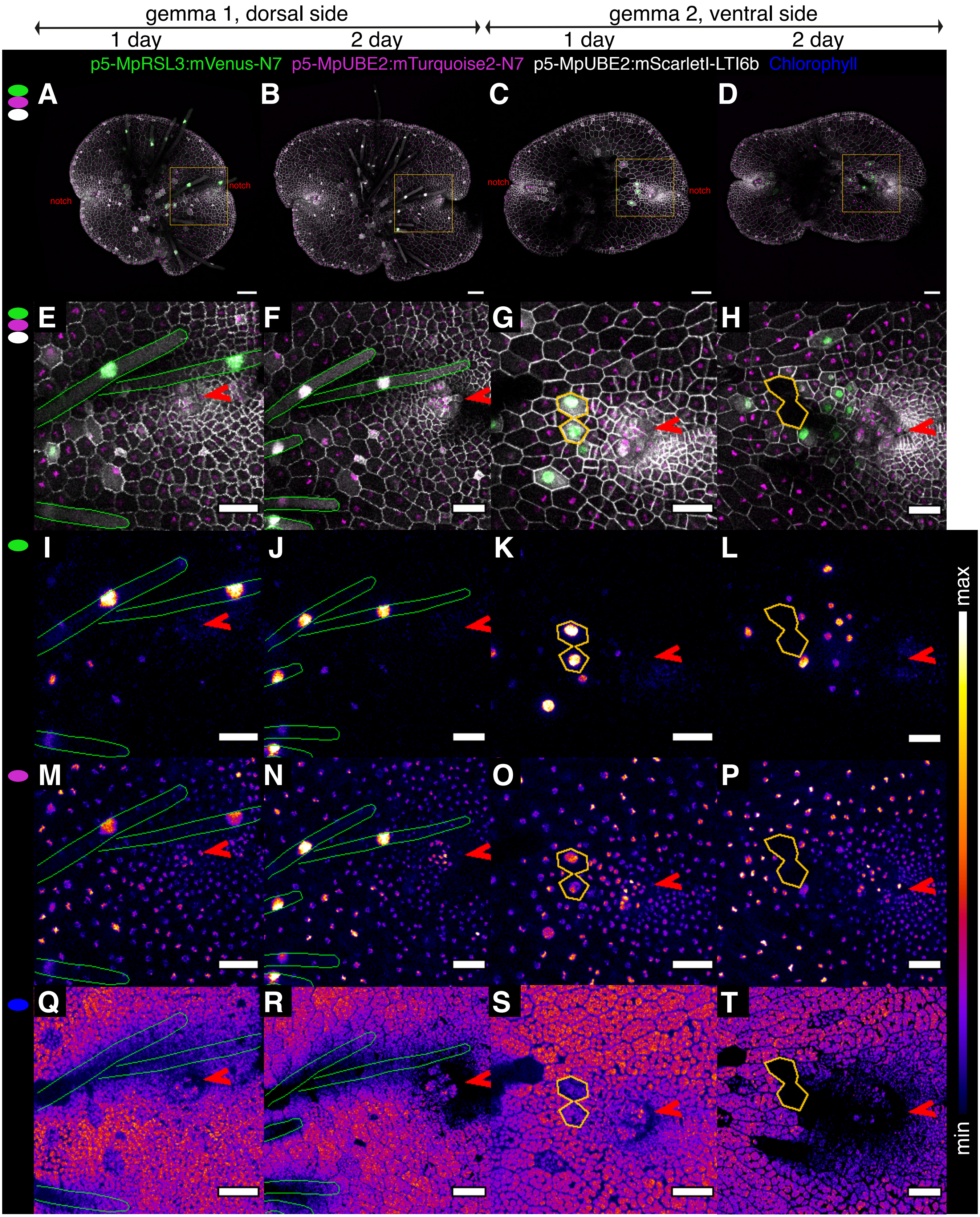

gemma 1, dorsal side
gemma 2, ventral side
1 day
2 day
1 day
2 day
p5-MpRSL3:mVenus-N7
p5-MpUBE2:mTurquoise2-N7
p5-MpUBE2:mScarletI-LTI6b
Chlorophyll
A
B
C
D
notch
notch
notch
notch
E
F
G
H
I
J
K
L
M
N
O
P
Q
R
S
T
max
min

**Figure S15.** Time course of MpRSL3 expression in early gemma development. Images of the dorsal side of gemmae, or the ventral side of a gemmae. In order to image the ventral side of gemmae, we grew plants on agar in an inverted plate (see Figure S14). Line expressing: p5-MpRSL3:mVenus-N7 (rhizoid marker, in green), p5-MpUBE2:mTurquoise2-N7 (constitutive nuclear expression, in magenta) and p5-MpUBE2:mScarletI-LTI6b (ubiquitous cell outline, in grey). Panels A-H show maximum intensity Z-axis projections gemmae surfaces with merged channels for mScarletI (grey), mTurquoise2 (magenta) and mVenus (green) channels. Image of whole gemmae (A-D) or a region (orange square in A-D and images E-T) from the center of the gemmae to one of the two notches (red arrow in E-T). At 1 and 2 days, the MpRSL3 marker can be detected in elongated rhizoids emerging from the central zone of the gemmae (A-D), like the ones delineated in green in the dorsal side images. In elongated rhizoids, the nuclei moves to the tip of the rhizoid cell (E,F). Moreover, only in the ventral side, the MpRSL3 marker is expressed in cells near the notch (G,H), which will become elongated rhizoids. Delineated in orange, a couple of cells expressing the MpRSL3 promoter at 1 day (G), which have already elongated at 2 days (with the nuclei above the Z-stack imaged and thus not in the image (H). (I-L). Levels of accumulation of the mVenus marker shown in false-color. (M-P) Levels of accumulation of the mTurquoise2 marker shown in false-color. (Q-T) Levels of chlorophyll autofluorescence shown in false-color. Scale bar: 100 $\mu m$ (A-D) or 50 $\mu m$ (E-T).

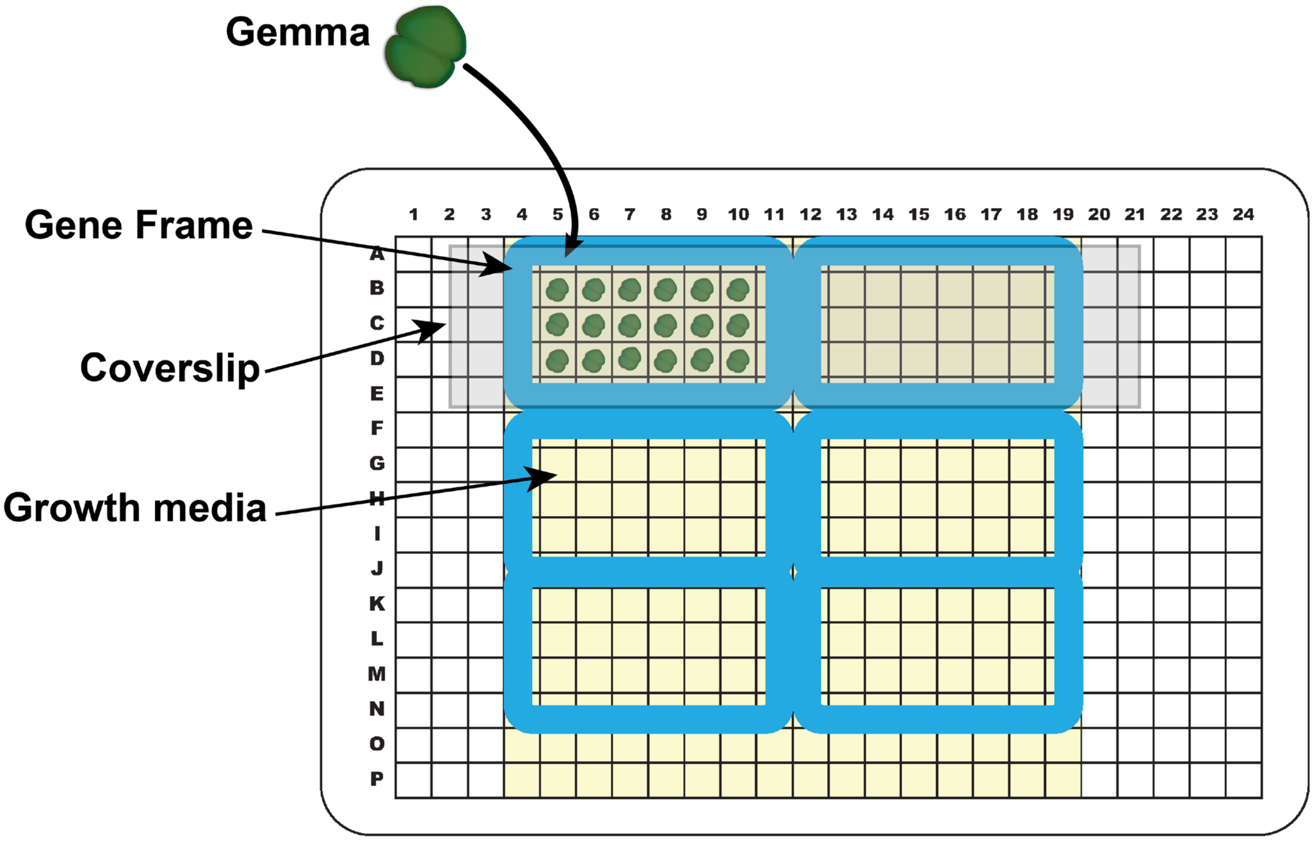


**Figure S16.** Culture of gemmae on multiwell plates for live imaging. Wells of a 384-well plate were filled with 0.5x Gamborg B5 media with 1.2% (w/v) agar. Single gemmae were placed at the center of each well. Gene frames and coverslips treated with anti-fog spray were used to cover and seal the wells. The setup allows automated imaging of samples.

**Table S1.** Quantification of CRISPR/Cas9 mediated editing of Mp*NOP1* using the OpenPlant toolkit CRISPR/Cas9 vectors

| | **Number of plants screened (L2_NOP1gRNA-Cas9-CsA)** | | | |
|---|---|---|---|---|
| | **total** | **wild type** | ***nop1*** | **chimeric** |
| **experiment 1** | **121** | **58** | **28** | **35** |
| | **%** | **47.93** | **23.14** | **28.92** |
| **experiment 2** | **133** | **69** | **22** | **42** |
| | **%** | **51.87** | **16.54** | **31.57** |

| | **Number of plants screened (L2_2xNOP1gRNA-Cas9-CsA )** | | | |
|---|---|---|---|---|
| | **total** | **wild type** | ***nop1*** | **chimeric** |
| **experiment 1** | **127** | **46** | **36** | **45** |
| | **%** | **38.01** | **29.75** | **37.19** |
| **experiment 2** | **125** | **61** | **25** | **39** |
| | **%** | **48.8** | **20** | **31.2** |

**Table S2**. List of primers used in this study

| | |
|---|---|
| | **For sequencing inserts in pUAP4 and pC vectors (pCk and pCs)** |
| UAP_F | CTCGAGTGCCACCTGACGTCTAAGAAAC |
| UAP_R | CGAGGAAGCCTGCATAACGCGAAGTAATC |
| pC_F | GCAACGCTCTGTCATCGTTAC |
| pC_R | GTAACTTAGGACTTGTGCGACATGTC |
| pC_R2 | CAATCTGCTCTGATGCCGCATAGTTAAG |
| | **For genotyping chloroplast transformants with construct I and III (Figure S10)** |
| P1_F | TTCGATTCCCGCTACCCGC |
| P2_R | GCTGGAACAGCTATCTTAACTTTTATTGGTGG |
| P3_F | CTAGCTTCTCCTGAAGGTTGGTC |
| P4_R | GTTGTCCCGCATTTGGTACAGC |
| | **For genotyping chloroplast transformants with construct II (Figure S10)** |
| P5_F | CGGAGACTAAAGCAGGTGTTGG |
| P6_R | ACAGATCCCATACTACCGCC |
| P7_F | CGGATCATGAAGTCAAGGGTTC |
| P8_R | GTTGTCCCGCATTTGGTACAGC |
| | **For cloning NOP1 gRNA1 and gRNA2 into L1-lacZgRNA-Ck2/3** |
| gRNA1-BbsI-F | CTCG-ATAGTCTTTGTGAGAGAATgt (modified from ref 2) |
| gRNA1-BbsI-R | TAAAac-ATTCTCTCACAAAGACTAT (modified from ref 2) |
| gRNA2-BbsI-F | CTCG-ATTAAGAGTGGAAGTTGCTTgt (modified from ref 3) |
| gRNA2-BbsI-R | TAAAac-AAGCAACTTCCACTCTTAAT (modified from ref 3) |
| | **For cloning NOP1 gRNA1 into L2-lacZgRNA-Cas9-CsA** |
| gRNA1-SapI-F | TCG-ATAGTCTTTGTGAGAGAATgt |
| gRNA1-SapI-R | AAAac-ATTCTCTCACAAAGACTAT |
| | **For genotyping *nop1* CRISPR mutants** |
| PR1_F | TGGGCCCATCTGTCACTAAG |
| PR2_R | CATGTCCACCAGTACGTG |
| PR3_R | GGTTGTATACGCCCGCTTTC |
